# Notochord and axial progenitor generation by timely BMP and NODAL inhibition during vertebrate trunk formation

**DOI:** 10.1101/2023.02.27.530267

**Authors:** Tiago Rito, Ashley R.G. Libby, Madeleine Demuth, James Briscoe

## Abstract

The formation of the vertebrate body involves the coordinated and progressive production of trunk tissues from progenitors located in the posterior of the embryo. In vitro models based on pluripotent stem cells (PSCs) replicate aspects of this process, but they lack some tissue components normally present in the trunk. Most strikingly, the notochord, a hallmark of chordates and the source of midline signals that pattern surrounding tissues, is absent from current models of human trunk formation. To investigate how trunk tissue is formed, we performed single-cell transcriptomic analysis of chick embryos. This delineated molecularly discrete progenitor populations, which we spatially locate in the embryo, compare across species, and relate to signalling activity. Guided by this map, we determined how differentiating human PSCs develop a stereotypical spatial organization of tissue types. We found that LATS1/2 repression of YAP activity, in conjunction with FGF-mediated MAPK activation, induced the transcription factor Bra/TBXT and facilitated WNT signaling. In addition, inhibiting a WNT-induced NODAL and BMP signaling cascade at the appropriate time regulated the proportions of different tissue types produced, including notochordal cells. We used this information to create an integrated 3D model of human gastrulation undergoing morphogenetic movements to produce elongated structures with a notochord and spatially patterned neural tissue formation. Together the data provide insight into the mechanisms responsible for the formation of the tissues that comprise the vertebrate trunk and pave the way for future studies of patterning in a tissue-like environment.

## Introduction

The formation of the vertebrate body axis is an evolutionary conserved process that requires the coordinated generation of multiple distinct cell types. This is evident in the trunk where the tissues that make up the body, such as the spinal cord, vertebral column, and gut are generated progressively from head to tail. A population of posteriorly located progenitors fuels this axis elongation by balancing self-renewal with differentiation into the tissues that comprise the trunk^1^. In all vertebrates, a central role is played by a structure known as the organiser or node, a morphologically recognisable cluster of cells located at the midline that regresses posteriorly as axial elongation progresses^2^. Cells emanating from the node give rise to midline tissues such as the notochord^3,4^, the mesodermal rod that provides mechanical and signalling cues to the embryo, and the floor plate, the ventral midline domain of the neural tube that patterns neural tissue. In addition, secreted factors from the node control and organise the fates of adjacent cells in the so-called node-streak border and primitive streak^5^. These axial progenitors go on to generate several trunk cell types including neural and mesoderm tissue.

The transient and dynamic nature of the node and adjacent progenitors has hindered a detailed understanding of how trunk tissues form. Although a variety of signals have been implicated, including WNT, BMP, NODAL and FGF signalling^6^, how these produce the diversity of cell fates necessary to form the body is not clear. The recent development of *in vitro* models of axis elongation provides a new experimental framework to investigate this question. These models are based on the differentiation of pluripotent stem cells exposed to combinations of agonists and/or antagonists of WNT, FGF, and/or TGFβ signaling. A variety of signalling regimes have been established that are thought to mimic the signalling conditions experienced by cells in and around the node. The resulting organoids have variously been termed gastruloids^7–10^, trunk-like structures^11^, SCOs^12^, or somitoids^13^/ axioloids^14^. Varying amounts of neural, endodermal, and mesodermal tissue have been reported in these different preparations. But despite the diversity of tissues generated and the remarkable self-organising capacity of these assemblies, several tissue components normally found in the vertebrate trunk are absent. Perhaps most notably the notochord, a defining feature of chordates, appears to be missing from most structures reported to date. Consequently, tissues that depend on signals from the notochord, such as the floor plate are also absent. The lack of notochordal cells from current models is surprising because embryological experiments suggest that notochordal progenitors are located close to the node, adjacent to cells that generate neural and somitic tissue^3,4^. This raises the possibility that the specification of notochord progenitors or the differentiation of notochord cells has specific signalling requirements, compared to other tissues, that have not been met in current models. To address this, we first set out to establish a more detailed understanding of the molecular identity and signalling mechanisms controlling axial progenitor pools and trunk formation.

## Results

### Single-cell transcriptomics of trunk development

To investigate the emergence of axial progenitor pools and the trunk tissue they generate, we took advantage of precise staging of chick embryos to select four closely timed stages, approximately 5-6h apart, with somite (S) numbers: 4S, 7S, 10S and 13S (HH8-11, Hamburger Hamilton Stages, **Figure 1a**). Tissue caudal to the third somite pair was dissected and the transcriptome of single cells analysed. To define cell types we employed fine-grained, unbiased Louvain clustering followed by aggregation into major cell populations based on marker gene expression (**Figure 1b, S1a**). The stages analysed encompassed the induction of HOXB/C9 (**Figure 1c, S1b**) and, in line with an axial elongation process continuously generating trunk tissues, the proportions of major cell types were remarkably stable across stages (**Figure 1d, S1c-d**). The largest differences detected between 4S and 13S were in the Intermediate/ Lateral Plate Mesoderm cluster (−7%) and in the neuromesodermal progenitors (NMPs) (+3 %), a transient axial progenitor population fated to both mesoderm and neural lineages (reviewed in ^15–17^), and PreNeural clusters (+3.4%). All stages combined, the dataset complements existing data from earlier developmental stages (HH4-HH7)^18,19^, and from tailbud and anterior portions of the embryo^20,21^.

**Figure 1.**
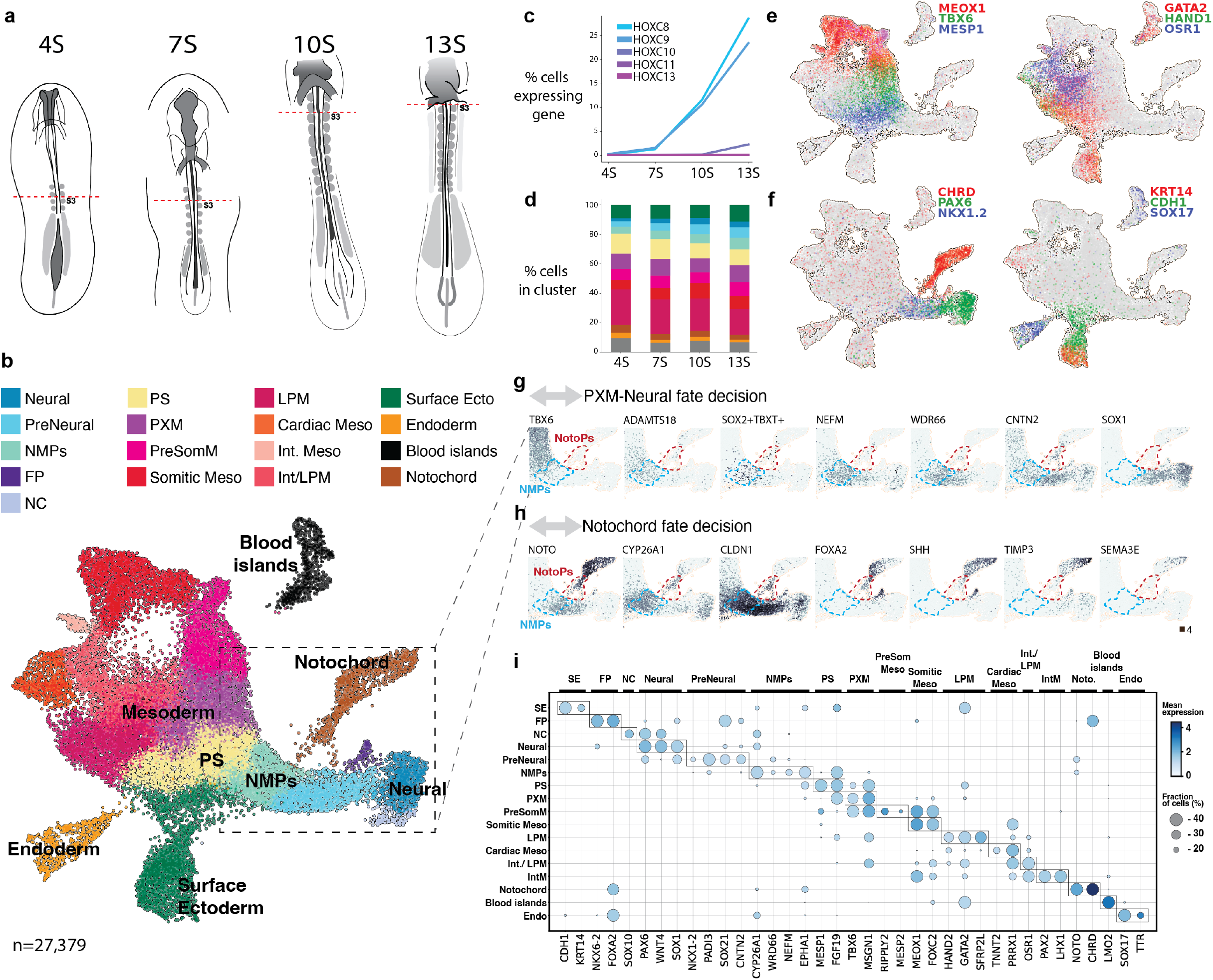
Single-cell transcriptomics of chick trunk development. **a,** Summary of chicken embryo dissections to generate single-cell RNA-seq datasets. Tissue caudal to the second somite pair in embryos with 4-Somites (4S), 7S, 10S and 13S (HH8-11) was dissociated. **b,** 2D embedding of single-cell data for all stages highlighting the different cell populations present during early vertebrate trunk formation. **c,** Proportions of cell types across different stages 4S-13S. **d,** Percentage of cells expressing different genes from the HOXC group showing induction of HOXC8-9 between 7-10S. **e-f,** Embedding of chick trunk (4-13S) coloured by different marker genes in the paraxial and lateral plate mesodermal compartment (e) and in notochord, neural, endoderm and surface ectoderm (f). **g-h,** Embedding detail plots showing the expression of several genes in the cell populations between paraxial mesoderm (PXM), neuromesodermal progenitors (NMPs) and Neural (g) and between NMPs and Notochord specification (h). Scale maximum counts is 4. **i,** Overview of gene expression levels and percentage of positive cells for marker genes used to distinguish cell types present in the trunk.

Analysis of gene expression identified cells of the expected tissues. MESP1, SNAI2 and MSGN1 mark primitive streak cells fated to somitic mesoderm. TBX6 was expressed in the paraxial and presomitic mesodermal cells, the latter with additional MEOX1 co-expression (**Figure 1e** *left*, **S1f**). Co-expression of MESP1 and RIPPLY2 identified cells at the so-called wavefront that are differentiating into the first somite (**Figure 1e** left purple, and **S1a** cluster 23) from TBX6+ pre-somitic mesoderm. Closely associated with the transition to paraxial mesoderm was ADAMTS18 (**Figure S1e**). This secreted metalloprotease cleaves fibronectin^22^ and promotes cell locomotion and epithelial-to-mesenchymal transition by decreasing cell adhesion, similar to other ADAMs in neural crest delamination ^23^.

Lateral plate mesoderm cells were identifiable by expression of HAND1/2, GATA2/4, CFC1, BMP4 and EVX1, it included a OSR1+ population, TBX20+TNNT2+ cardiac mesoderm and PAX2+LHX1+MNX1+ intermediate mesoderm (**Figure 1e *right* and S1f)**. Notochord cells expressed high TBXT, NOTO, SHH and CHRD (**Figure 1f *left*** and **S1g**). SHH also marked the floor plate and SOX17+ endodermal cells, the latter included a MNX1+HOXB1+ subcluster of the pancreatic lineage (**Figure S1f-g**).

A CDH1+ surface ectoderm population was composed of more mature surface ectoderm expressing KRT14/17, TFAP2A, DLX5, EPHA1 and WNT6 and a group of cells co-expressing MESP1 and CDH1 presumably marking cells at the embryo’s surface near the primitive streak^24^ (**Figure 1f** *right* and **S2a)**. PAX6 expressing neural cells (**Figure 1f** *left*) included ventral (NKX6-1/ 2+) and dorsal (PAX3+MSX1+) progenitors and SOX10 expressing neural crest cells (**Figure S2b**). In addition, a population of PreNeural cells expressing NKX1-2 (Sax1) was evident^25,26^. The data revealed previously uncharacterised markers of this population such as PADI3 and CA2, and the contactin CNTN2, a cell adhesion/recognition molecule from the immunoglobulin superfamily. A recent single-cell study of early anterior neural induction in chick^21^ identified common genes, including ZIC2 (early PreNeural) and MAFA (late PreNeural/Neural) as well as others such as GLI2, RFX3, ZNF423 and TAF1A that were more broadly expressed at these stages (**Figure S2c**).

We mined the data to identify transcriptomic signatures of axial progenitor populations involved in generating trunk tissue. Two progenitor populations stood out: NMPs and putative notochord progenitors. The NMP cluster included the highest density of SOX2+TBXT+ cells, markers typically used to identify these progenitors (**Figures 1g** and **S2d**), as well as enrichment in other, more broadly expressed genes that were previously associated with NMPs such as NKX1-2 ^27^, EPHA1^28^, CDX2 and CYP26A1 ^29^. Low levels of NOTO were detected in the NMPs in agreement with previous observations in chick^30^. Expression of CLDN1 and F2RL1 (Par2, **Figure S2d**) were also present in this population, emphasizing the proposed epithelial nature of NMPs ^4,31^. Additionally, NEFM, an intermediate filament, and WDR66 (ENSGALG00000004365 in chick), a cilia and flagella associated protein, were also expressed in NMPs. By contrast, NOTO+CHRD+ notochord cells contained at least two distinct populations: one which we hypothesized to correspond to the posterior node, i.e. the cell population around the median pit (**Figure 1h** - NotoPs), and the second that appeared to be more mature notochord. The former additionally expresses FOXA2, CDX2, CYP26A1, CLDN1, CNTN2 and GNOT2, whilst the more mature notochord cells express SHH, NOG, LEFTY and also TIMP3, a metallopeptidase inhibitor, and SEMA3C/E, a semaphorin implicated in control of cell morphology and motility.

Near identical cell populations can be found by analysis of existing mouse (E8.0-E8.5)^32^ and macaque (CS9-CS11)^33^ single-cell transcriptomic data highlighting the remarkable conservation of trunk formation across vertebrates (**Figure S3a-c**). However, the number of notochord cells in mouse were much lower than in chick and very few progenitors were captured, perhaps due to the small notochord size in mouse. The macaque data showed a similar, close relationship between NMPs and notochord cells as in chick, marked by an intermediate cell population expressing FOXA2, NOTO and CDX2. Crucially, the NMP cells, as in chick, exhibited the highest proportion of double-positive Sox2+T+ cells and expressed Cyp26a1 and Nefm. Together, we detected both established and new gene markers for each population (**Figure 1i, Table S1**).

### Cytoskeletal components and TGFβ inhibitors delineate axial progenitor populations

To spatially map progenitor populations defined in the transcriptome analysis in chicken embryos we used third generation multiplexed quantitative RNA in situ hybridization (HCR)^34^. NEFM, which was expressed in NMPs, marked the dorsal part of the node-streak border where the SOX2+TBXT+ NMPs reside^35^, extending from the anterior primitive streak to the lateral edge of the pre-neural domain around the caudal node (**Figure 2a**). The equivalent domain in mouse has a smaller inverted-U shape surrounding the primitive streak reflecting the distinct geometry of rodent embryos^16^. NEFM was also found further away from this region in the caudal portion of somites and at the folding edges of the neural tube (**Figure S4a**). CNTN2, which delineates PreNeural cells, marked a complementary domain rostral to the node. Similar to NEFM, CNTN2 was mostly dorsal but we noted expression in some ventral cells near the caudal onset of NOTO. Rostrally, NOTO and CNTN2 form two mutually exclusive dorsal-ventral domains before any morphological segregation between notochord and neural plate was apparent (**Figure 2b**). CNTN2 expression also spatially preceded the more rostral expression of the neural marker SOX1 (**Figure S4b**). CYP26A1 and ADAMTS18 were expressed more caudally in a narrow ellipsoid domain abutting CNTN2. This domain partially overlapped NEFM but extended ventrally from the primitive streak to just above the medial pit (**Figure S4c**).

**Figure 2.**
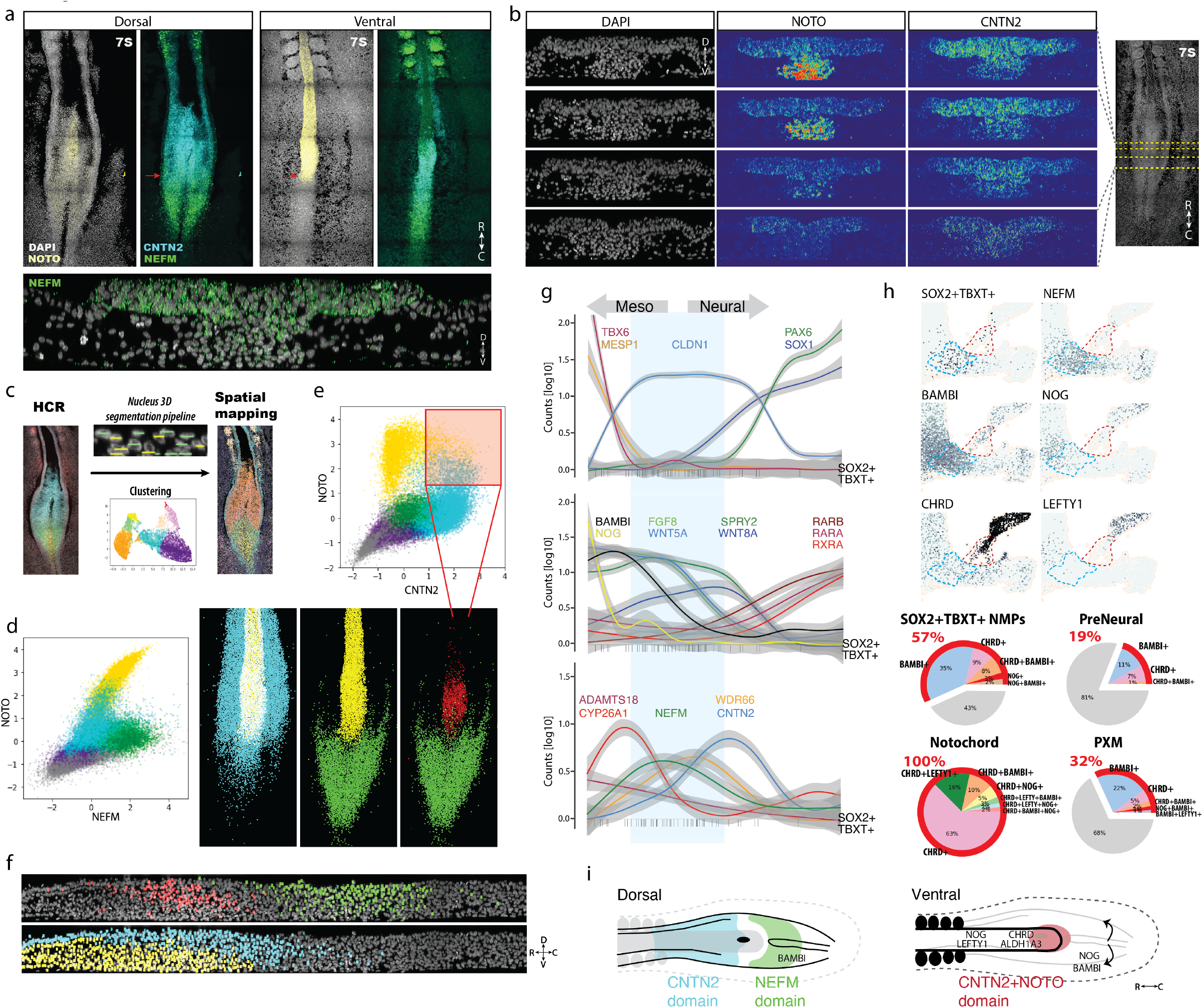
Axial progenitors delimited by cytoskeletal components and TGFβ inhibitors. **a,** Multiplexed RNA fluorescence in situ hybridization (HCR) image of a 7S chick embryo stained for NOTO in yellow, NEFM in green and CNTN2 in cyan. **b,** optical section of an HCR image showing dorso-ventral segregation of NOTO and CNTN2 caudal of the median pit. **c,** single-cell quantification of HCR signal using custom Nucleus 3D segmentation pipeline. **d,** Scatter plot of HCR signals per single cell for NOTO and NEFM in 7S chicken embryos; colouring reflects hierarchical clustering for K=5. Clusters were spatially mapped back to the embryo. **e,** Cells with high NOTO and high CNTN2 HCR signals map to the medial pit of the 7S embryo (red cells in d). **f,** Axial optical section of 7S embryo showing dorso-ventral organization of the different HCR clusters. **g,** Transcriptional similarity ordering (pseudotime analysis) of chick single cells from paraxial mesoderm, NMPs and Neural clusters showing the expression of CNTN2, NEFM relative to other lineage and signalling genes. Vertical black lines in the bottom of the plots show the position in the pseudotime axis of cells co-expressing both SOX2 and TBXT. **h,** Embedding detail with gene expression of TGFβ inhibitors relative to NMPs. Pie charts with percentage of cells (red) in each cell population expressing at least one BMP or NODAL inhibitor. **i,** Schematic of the dorsal and ventral view of an elongating 7S embryo. Colour highlights the HCR domains identified with cytoskeletal and adhesion components, and the expression of key TGFβ inhibitors.

Using a custom imaging analysis pipeline, *Nucleus* (see Material and Methods, **Figure S5**) we clustered cells based on HCR signals (**Figure 2c**) and mapped these to the embryo (**Figure 2d**). We identified cells in the median pit of the posterior node and found that these correspond to the intersection of high NOTO and high CNTN2 expression (**Figure 2e-f** red **and S6a**). The transcriptome data indicated that these cells corresponded to the early notochord progenitors and expressed CLDN1 along with CYP26A1 and FOXA2. We confirmed FOXA2 expression in the same spatial region by immunostaining (**Figure S6b**). Together, the spatial analysis and transcriptome data delineate specific domains within the posterior growth zone corresponding to different progenitor populations.

Complex signalling networks regulate the behaviour of cells in the node-streak border^6,16^. Using pseudotime-sorting^36^ to construct a fate decision axis reflecting both time and space indicated a large CLDN1+ cell population between SOX1+ neural progenitors and MESP1+TBX6+ PXM that included NEFM expressing NMPs (**Figure 2g**). Characteristic dynamics of components of the WNT, FGF and RA signalling pathways were evident along this trajectory^6,37^. For instance, along the trajectory to neural identity, WNT5A and FGF8 were first downregulated, then WNT8A and SPRY2, followed by the up-regulation of WNT4, RXRA and RARB retinoic acid receptors. An equivalent analysis conducted using mouse data showed the same overall behaviour despite differences in some specific gene orthologues (**Figure S6c**). Consistent with prior studies^38,39^, multiple BMP and NODAL inhibitors were expressed. NMPs expressed high levels of BAMBI – a pseudoreceptor inhibiting BMP and NODAL signalling^40^, mirrored in mouse by FST; and the underlying populations of nascent paraxial mesoderm and notochord have a high percentage of cells expressing NOG and CHRD (**Figure 2g-h**). Additionally, the NODAL inhibitor LEFTY1 was expressed in mature notochord cells.

Together, these results indicate that rather than a field of equivalent or spatially heterogeneous progenitors there is a highly structured architecture in and around the caudal node. The metalloprotease ADAMTS18 and CYP26A1 mark ingressing cells at the caudal primitive streak extending ventrally past the node. Dorsally, NEFM together with CYP26A1 and BAMBI mark epithelial neuromesodermal axial progenitors. In the rostral direction, a small ventral domain of FOXA2, CNTN2 and NOTO demarcate the median pit expressing BMP inhibitors, whilst in the preneural tube, dorsal CNTN2 apposes ventral NOTO before the first SOX1+ cells mark the future neural tube (**Figure 2i**). Combining the single cell transcriptome data with *in vivo* mapping provides new insight into the organisation of fate transitions driving axis elongation and the spatial dynamics of the signalling pathways controlling them.

### A minimal *in vitro* model of posterior axial fate decisions

The spatial organization in the embryo prompted us to develop a system in which to test the function of specific signalling pathways in the specification of axial progenitors and the formation of trunk tissue. Taking advantage of the evolutionary conservation of trunk cell populations, we established an *in vitro* model using human embryonic stem cells. Several *in vitro* protocols have been proposed to generate axial progenitors ^41–44^ (reviewed in ^17^). Most of these rely on WNT and FGF activation and we used this as a starting point to generate human trunk progenitors.

We first employed a 3-day monolayer protocol consisting of exposure to FGF2 and CHIR99021 (CHIR), a WNT agonist acting via GSK3beta inhibition. Reflecting our observations in the chick, we added BMP and NODAL inhibition. This resulted in a high frequency (89%, n=6,997) of double positive SOX2+TBXT+ cells (**Figure S7a**). However, we observed disorganised and heterogenous levels of these two markers. We asked if geometric confinement might provide a more ordered pattern of gene expression^45,46^. Applying the same treatment to hESCs seeded on circular micropatterned laminin substrates resulted in a striking pattern of cell fate organisation at day 3: SOX2-high cells were in the centre and TBXT-high cells at the edge of the colony (**Figure 3a, Movie 1**). Clumps of TBXT expressing cells formed morphologically distinct structures at the periphery of colonies and the number of these aggregates scaled with the diameter of the micropattern: the smallest diameter colonies (200um) typically contained a single TBXT aggregate whereas the larger 500um diameter colonies had 2-3 clusters (**Figure 3b**). We term these cellular assemblies posterior neuruloids.

**Figure 3.**
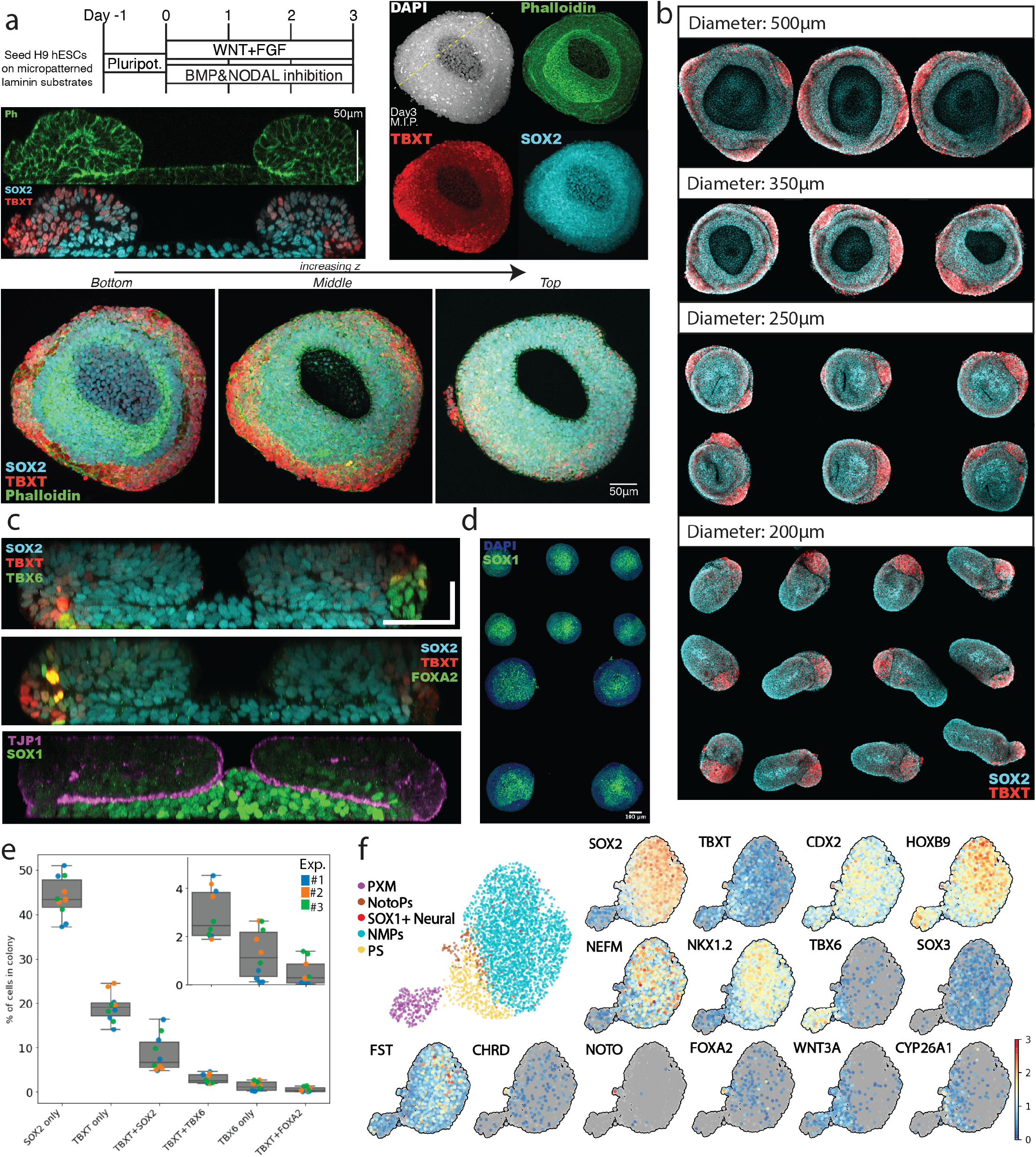
*In vitro* model of human trunk formation. **a,** Summary of the posterior neuruloid platform using colonies of human embryonic stem cells grown on micropatterned laminin substrates. The protocol employs a 3-day treatment with WNT-agonist (CHIR99021) and FGF2 together with BMP and NODAL inhibitors (SB431542 and LDN193189). Maximum intensity projection (M.I.P.), optical section and single-z plane images of immunofluorescence staining of SOX2 (neural), TBXT (mesodermal) and phalloidin (F-actin). **b,** Size-dependant organization of the posterior neuruloid with clustering of TBXT+ mesodermal tissue (red) for colony diameters between 200-500μm. **c,** Medial optical sections of immunofluorescence showing the location of paraxial (TBX6) and axial (FOXA2) mesoderm; SOX2 and the SOX1 neural marker, as well as the tight-junctions marker ZO-1 (TJP1). **d,** SOX1 immunostaining overview illustrating the robustness of the platform. **e,** Nucleus 3D quantification of ~8,750 cells in each posterior neuruloid colony showing the proportions of cell types obtained by immunofluorescence. **f,** Single-cell RNA-seq characterization of posterior neuruloids. 2D embedding showing the different cell populations and the expression of the main marker genes distinguishing them.

We assayed additional markers to assess cell type identity in neuruloids. Cells in the peripheral aggregates expressed the paraxial mesodermal marker TBX6. Reminiscent of mouse ESCs differentiations^47^, a few TBXT high cells co-stained with FOXA2 indicating the presence of node-like cells or notochord progenitors. The central SOX2-high cells expressed the neural marker SOX1 (**Figure 3c**) and this neural induction was reproducible across colonies (**Figure 3d**) and multiple experiments. Assaying TJP1 (ZO-1) indicated that the cells had acquired a coordinated apical-basal polarity.

To document how neuruloids formed we used timelapse imaging. The colonies were highly dynamic and had a stereotypical inward movement driven by cells located in an intermediate ring between the edge and the colony’s center (**Movie 2**). We estimated this movement to be ~4um/hour and this caused the folding of the apical side towards the centre (**Figure 3c** bottom, ZO-1 stain) resulting in the typical donut-shape structure seen in colonies at Day 3.

To quantify cell numbers and identities, we trained and applied our *Nucleus* 3D segmentation tool using images of densely packed nuclei in micropatterns. The average number of cells per colony was 8749 (SD=1521, N_experiments=3, n_colonies=3). The proportion of fates was largely consistent across different experiments with 40-50% of cells positive for SOX2 only, 20% for TBXT only and 10% double-positive. Notochord TBXT+FOXA2+ progenitors and TBX6+TBXT-cells formed a minority (2%, **Figure 3e**). Of note, notochord cells also represent 3-5% of the captured cells in the chick datasets. We subdivided the intensity levels of SOX2 and TBXT expressing cells into three categories. Cells expressing high TBXT or high SOX2 cells were located at the edge and centre of the colony, respectively, while cells with mid-levels of both factors were located on the top of the colony, in an intermediate ring position closer to the edge (**Figure S7b**).

Finally, to allow comparisons with chick and other vertebrates, we performed single-cell RNA-seq. These data confirmed the posterior character of the colonies by expression of CDX2 and HOXB9 and presence of PXM and PS-like populations expressing WNT3A, DKK1, MIXL1, TBX6 and FOXC2 (**Figure 3f and S7c**). A large fraction of cells expressed both SOX2 and TBXT mRNA. As in the chick data, this NMP cluster expressed NEFM (66% of cells in cluster, n=1739/ 2623). CYP26A1 was also present however was more restricted to PS cells. The expression of NOTO, CHRD and SHH, in addition to FOXA2 and TBXT in a group of cells confirmed the presence of notochord. Taken together, the data indicate the presence of paraxial mesoderm, notochord and neural progenitors, cell populations present in the caudal embryo during neurulation stages, suggesting that posterior neuruloids offer a good model to investigate the mechanisms of trunk formation.

### YAP inactivation and persistent pERK1/2 modulate the WNT response to regulate TBXT

To investigate the relationship between the signalling pathways identified in axial progenitors in the chick embryo and the pattern of cell types is established in posterior neuruloids, we examined early stages of their development. We looked for evidence of FGF-MAPK pathway activation. Immunostaining for phosphorylated ERK1/2 (pERK1/2), a target of MEK1/2, showed colony-wide activation as early as 3h after addition of the induction media. By 12h, pERK1/2 was restricted to a ring at the edge of the pattern and remained so at 24h. Only a few cells in the centre of the colony contained pERK1/2. This pattern was superseded by TBXT expression. At 12h TBXT+ cells were found in a salt-and-pepper fashion throughout the colony and by 24h TBXT expression had become localized to the edge of colonies (**Figure 4a**). These dynamics were reproducible across colonies and experiments (**Figure S8a**). The pERK1/2 TBXT-expressing ring was approximately 2-3 cell diameters wide. Moreover, application of the MEK1/2 inhibitor PD98059 abolished the ring signature indicating that both pERK1/2 and TBXT expression were dependent on FGF signalling (**Figure 4b**, **S8b**). Furthermore, increasing FGF concentrations did not significantly change the width of the pERK1/2 ring at 24h, despite a progressively larger TBX6+ domain at the periphery by day 3 (**Figure S8c**). In the absence of FGF, no TBX6+ cells were present at day 3, indicating endogenous FGF signalling was insufficient to drive paraxial mesoderm formation. The pERK1/2 ring was also significantly diminished in the absence of the WNT agonist, indicating synergy between WNT and FGF pathways (**Figure 4b, S8b**). By 48h, pERK1/2 was still detected at the edge where the first TBX6+SNAIL+ cells were induced (**Figure 4c**). These cells constitute a delaminating population that ingresses underneath the initially flat epithelial layer of the micropattern to initiate the 3D, donut-like shape of the posterior neuruloid, reminiscent of the behaviour of prospective mesodermal cells ingressing at the primitive streak.

**Figure 4.**
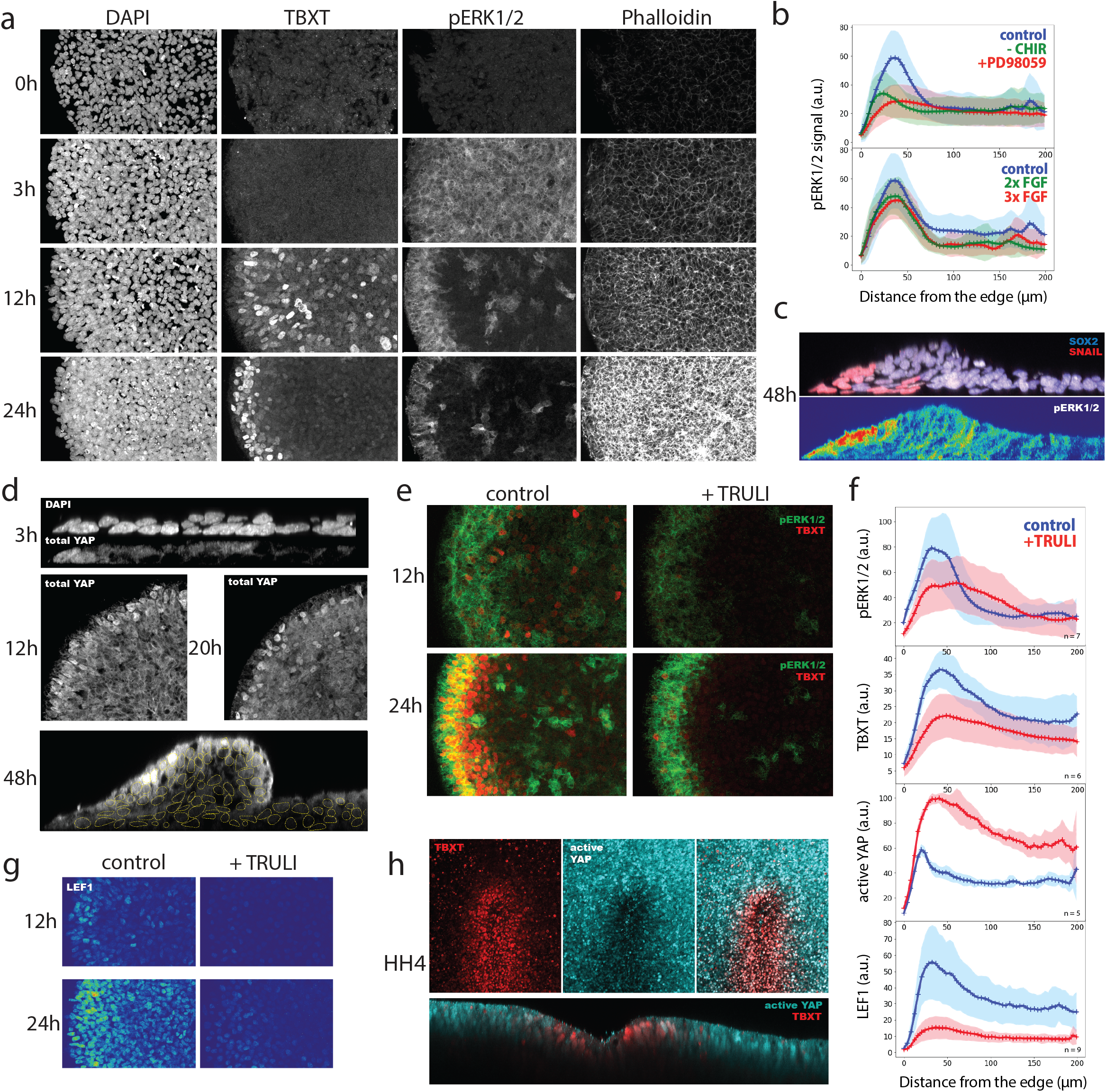
Induction of TBXT by coordinated FGF, WNT and Hippo signalling. **a,** Time-course of immunofluorescence for TBXT and pERK1/2 during the first 24h of the posterior neuruloid. **b,** Quantification of radial pERK1/2 levels at 12 hours post-induction (N=2, n=3) without the WNT-agonist (CHIR) and with varying concentrations of FGF and its inhibitor (PD98059). **c,** Immunofluorescence for pERK1/2 at 48 hours post-induction showing that high levels co-localize with SNAIL and TBX6 induction at the edge of the colony. **d,** Localization of total YAP during posterior neuruloid formation. **e,** pERK1/2 and TBXT at 12 and 24 hours in the presence of TRULI, a LATS1/2 inhibitor, causing drastic reduction in TBXT positive cells. **f,** Quantification of radial levels of several markers at 12 hours post-induction of the posterior neuruloid and in the presence of an activator (TRULI) of YAP-mediated signalling. **g,** Immunofluorescence staining of LEF1 showing reduced signalling in the presence of TRULI. **h,** Staining of TBXT and active YAP (unphosphorylated) in a chicken embryo at full primitive streak stages (HH4).

To explore the induction of TBXT, a target of WNT^48^, we examined YAP signalling as the pathway has been associated with modifying WNT signalling ^49,50^ and with sensing cell-cell mechanics via LATS1/2 and MST1/2^51^. The role of YAP signalling during trunk formation remains unclear despite YAP null mice exhibiting a truncated axis^52^ and YAP being implicated in controlling the segmentation clock^53^. As previously observed in micropatterned hESCs ^54^, we found that cells at the periphery retain nuclear YAP whilst internal cells progressively exclude YAP from the nucleus during the first 24h (**Figure 4d**). To test the relationship between YAP and TBXT at the edge we perturbed YAP signalling. A LATS1/2 inhibitor, TRULI^55^, which promotes nuclear YAP accumulation (**Figure S9a**), caused a marked reduction in TBXT+ cells at 24h (**Figure 4e**), leading to the absence of axial and paraxial mesodermal derivatives at day 3 (**Figure S9b-c**). By contrast, the ring of pERK1/2 at 24h was relatively unaffected albeit with lower levels. This decrease in pERK1/2 was also observed at 12h (**Figure 4f and S9d**). Thus, the ring of pERK1/2 activity is not dependent on YAP, although YAP signalling can modulate pERK1/2 levels. By contrast TBXT expression is inhibited by YAP signalling. This led us to investigate the relationship between YAP and WNT signalling.

YAP activation has been linked to promoting^56^ and to suppressing^57^ WNT signalling. To investigate how YAP activation impairs TBXT induction we used LEF1 expression as readout of WNT signalling^58^. As early as 12h, the control pattern had mounted a WNT response which was higher at the colony’s edge with levels of LEF1 decreasing toward the centre (**Figure 4f-g**). In TRULI treated samples expression of LEF1 was greatly diminished or undetectable at 12h, suggesting that nuclear YAP inhibits WNT signalling activity. At 24h LEF1 was observed in TRULI treated colonies but compared to the control, there were fewer positive cells and lower levels, pointing to YAP activity delaying and weakening the WNT response. This suggests a YAP activation blocks TBXT induction by inhibiting WNT signalling. To investigate if the absence of YAP activity also correlated with TBXT expression *in vivo*, we assayed HH4 chicken embryos for active YAP. Indeed, TBXT expression in cells of the primitive streak is accompanied by downregulation of active, unphosphorylated YAP (**Figure 4h**).

Together, these data suggest a stepwise model leading to the stereotypical spatial organization of the posterior neuruloid: WNT and MAPK activation result in TBXT and pERK1/2 induction facilitated by LATS1/2 activity and nuclear YAP exclusion. Persistent WNT and FGF signalling at the edge stabilize TBXT expression and result in the upregulation of TBX6 and SNAI2.

### Timing of BMP and NODAL inhibition orchestrates self-organization of caudal cell fates and can be harnessed to generate a notochord-like population in gastruloids

Next, we turned our attention to the small number of notochord-like cells in posterior neuruloids and sought to identify signals responsible for inducing these cells. Cross-species comparisons identified a population of TBXT+FOXA2+CDX2+ cells, most evident in chick and macaque. These cells expressed NOTO and CHRD together with FGF and WNT ligands and occupied a region of gene expression space between NMPs and mature notochord (**Figure S10a**). The expression of several BMP and NODAL inhibitors in cell populations in and around the node (**Figure 2**) prompted us to test whether altering the timing and duration of TGFβ signalling inhibition affected the cell composition of posterior neuruloids (**Figure 5a**).

**Figure 5.**
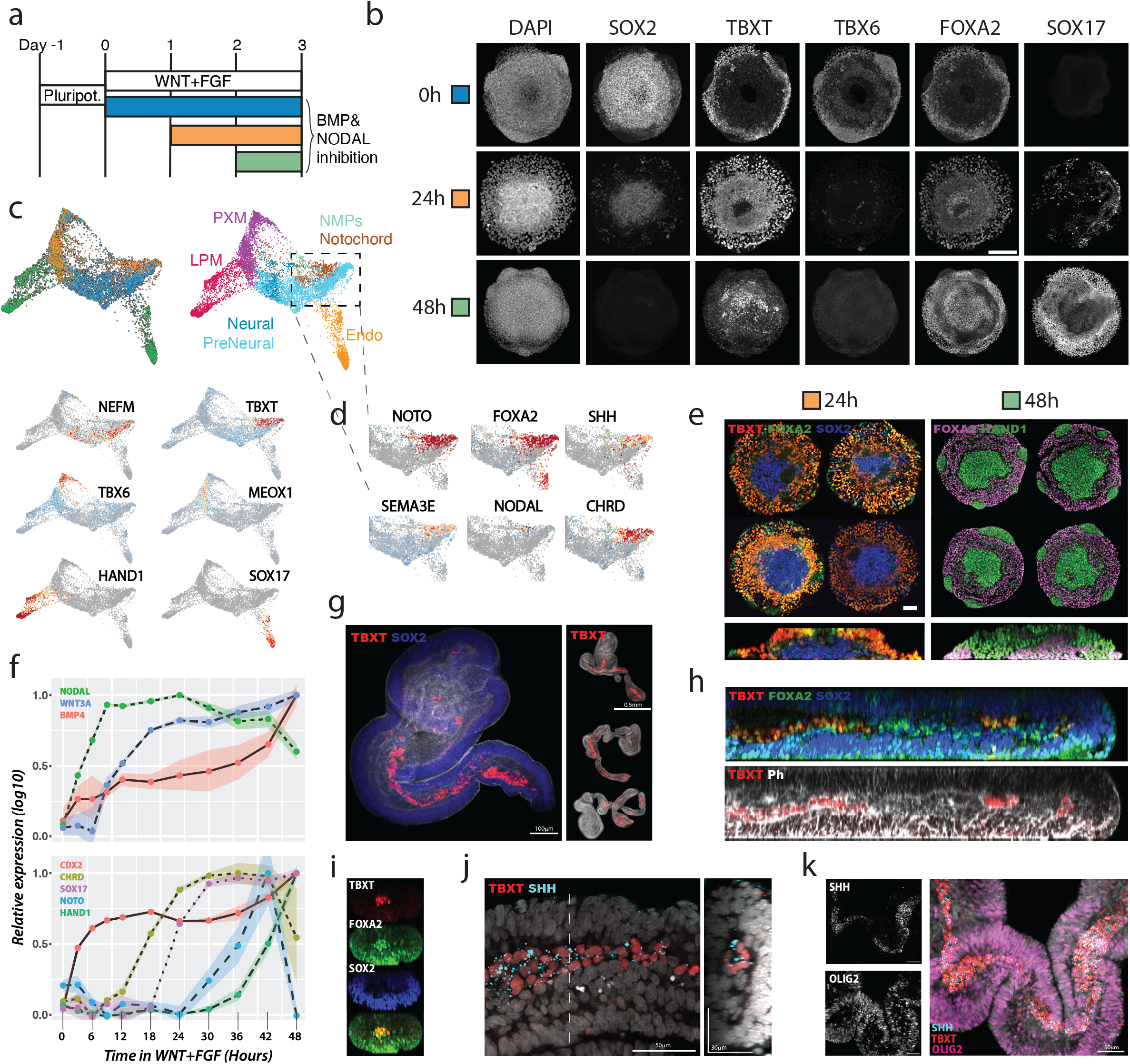
Timing of BMP and NODAL inhibition and the generation of notochord-like cells. **a,** Schema for posterior neuruloid differentiation with modifications to the timing and duration of BMP and Nodal inhibition. The control condition with constant inhibition is depicted in blue (0 hours); 24 hours delayed inhibition in orange; and inhibition starting 48 hours after induction in blue. **b,** Immunofluorescence for cell type markers (SOX2, TBXT, TBX6, FOXA2, SOX17) for the different BMP and NODAL delay treatments. Scale bar 100μm. **c,** single-cell RNA-seq characterization of the different inhibition treatments. Combined embedding showing largely non-overlapping cell populations generated in the different treatments. Colouring reflects the timing of BMP and NODAL inhibition, *i.e*. the different treatments (*top left plot*), the cell lineages generated (*top right plot*) and the expression of a few key genes of each population (*bottom plots*). **d,** Embedding detail showing the expression of notochord markers in the TBXT-high region of the plot, mostly cells from the 24h-delay treatment. **e,** Reproducibility and spatial segregation of the different cell populations present in the 24h- and 48h-delay treatments. Scale bar 50μm. **f,** Time-course of relative expression of morphogens and cell fate markers from qPCR data collected every 3-6h during the first 48h of treatment with WNT agonist (CHIR) and FGF. **g,** 3D gastruloid protocol combining 24h delay treatment with additional 4 days in the presence of a retinoid derivative generates elongated structures with a TBXT expressing cells internally surrounded by SOX2+ epithelial cells. **h-i.** Optical sections highlighting TBXT and FOXA2 co-localization. **j,** Immunofluorescence showing SHH accumulation in the TBXT+ notochordal cells. **k,** OLIG2 staining showing ventral neural patterning in the cells surrounding the notochord.

Almost no SOX2+ or TBXT+ cells were produced in the absence of BMP and NODAL inhibition or by delaying inhibition until 48h after addition of WNT and FGF, indicating that TGFβ signaling inhibition is necessary for the specification of NMP cells (**Figure 5b**). Instead, a dramatic increase in endoderm cells, expressing both SOX17 and FOXA2 was evident. Moreover, single-cell RNA-seq of these conditions revealed the presence of another population of cells expressing low TBXT, CDX2, BMP4, HAND1/2 and GATA3/4/6, which is consistent with lateral plate mesoderm (**Figure 5c, S10b)**. Strikingly, the different populations were organized in a stereotypical fashion indicating not only on-going cell fate specification but also distinct mechanical or cell adhesion cues driving morphogenetic behaviour (**Figure 5e**).

A 24h delay to the addition of TGFβ inhibitors also resulted in a marked reduction of SOX2+TBXT+ cells. In this case, however, many cells adopted a TBXT+FOXA2+ fate, suggesting a notochord identity. These cells were located on the colonies’ surface and periphery whereas cells in the centre were SOX2+ (**Figure 5b**). Single-cell RNA-seq data confirmed the cell lineages generated by the different delays versus on-going inhibition (**Figure 5c**). Importantly, the TBXT+FOXA2+ cells co-expressed NOTO, SHH, CHRD and SEMA3E, similar to the notochord population in chick (**Figure 5d**). In addition, SOX2 expressing cells were apparent along with a few paraxial mesoderm and pre-somitic cells, which expressed TBX6 and MEOX1. To validate the assignment of cell identity, a classifier trained on chick and mouse cell clusters correctly placed the different neuruloid populations in our assigned cell fates (**Figure S10c-d**). Thus, transient TGFβ signalling results in the generation of notochord-like cells along with neural and paraxial mesoderm, whereas prolonged signalling generates endodermal and lateral plate mesoderm tissue.

The effects of delaying TGFβ signalling on colony formation prompted us to examine the dynamics of morphogen gene expression (**Figure 5f**). We found that the WNT+FGF signalling regime rapidly induced NODAL during the first 9h, before the induction of WNT3A ligand itself (12-18h). CHRD expression preceded SOX17 (24-30h) by about 6h and, although NOTO is up-regulated at 42h, it rapidly decreases as BMP4 and HAND1 are up-regulated. These results not only argue for a limited window in which NODAL and BMP signalling specifies notochord fate but also highlight the need for on-going TGFβ inhibition to maintain NMPs.

Next we sought to adapt the signalling conditions defined on micropatterns to 3D culture. Current 3D *in vitro* models of human gastrulation feature NMP-driven axis elongation and the generation of somitic and neural tissue with a dorsal identity, but they lack a defined notochord population and generally lack ventral somitic and neural cell types, unless SHH signalling agonists are also included^9,59,60^. To test whether timed inhibition of TGFβ signalling was sufficient to promote the production of notochord cells in 3D models, we exposed aggregates of hESCs to the same signalling regime that generated notochordal cells in micropatterned culture (**Figure 5a** orange) then cultured these for four additional days in the presence of a retinoic acid precursor (**Figure S11a**). This resulted in the elongation of the aggregates (**Figure 5g, S11b**), 75% (n=36) of which featured a stripe of TBXT+ cells in their interior that was suggestive of the presence of a notochord (**Figure 5g, S11c**). The elongation was present in all experiments (**Figure S11d**). The outer cell layer was SOX2+ and had an epithelium-like morphology characteristic of a neural character. The expression of FOXC2+ cells in the interior of the tubular 3D structure also pointed to the generation of somitic tissue (**Figure S11e**). Consistent with the presence of notochord-like structures, TBXT+ cells expressed FOXA2 and were SOX2 negative while surrounding SOX2+ cells were TBXT- (**Figure 5h-i**). Moreover, we detected SHH expression in the TBXT+ streaks indicating that patterning of neural tissue could be occurring (**Figure 5j**). Indeed, we detected FOXA2+SOX2+ cells, OLIG2, and NKX6.1 expression indicating the presence of floor plate and motor neuron progenitors (**Figure 5k, S11f-g**). Ventralization of neural tissue was closely associated with the presence of notochord-like cells (**Figure S11f**). The generation of these cell types emphasises the role of dynamic signalling interactions within the node-streak border; further understanding these time-dependent relationships will enhance our ability to study cell fate choices.

## Discussion

In this study we identify and locate the major axial progenitor populations orchestrating vertebrate trunk formation by analysing the transcriptome of closely staged chicken embryos – a workhorse for functional studies for which single-cell genomic data has been scarce. Comparison with data from mouse and macaque embryos and an *in vitro* model of human trunk formation identified analogous progenitor populations, consistent with the highly conserved nature of vertebrate trunk formation. The chick data highlighted genes with potential functions in these progenitor pools and provided insight into the spatial and relational organisation of the cells fuelling axis elongation. The spatial and transcriptional proximity of notochord and axial progenitors revealed by the analysis suggests a close relationship between these cell types, consistent with prior embryological studies^3,4^.

Guided by the detailed *in vivo* map we used micropatterned hESCs colonies to develop novel protocols for generating axial progenitors and trunk tissue. This proved to be an attractive platform to study trunk tissue formation, offering a trade-off between complexity and tractability, and allowed us to define conditions in which neural, mesoderm and notochord progenitors were juxtaposed in single colonies. This complements existing models of gastrulation employing BMP4^45^ and expand primitive streak models that use WNT3A ligand 61-6^3^. We showed that despite generalized WNT activation, LEF1 and TBXT induction was restricted to the edge of micropatterns. This required persistent MAP kinase pathway activity and was facilitated by YAP inactivation, which appears to enhance WNT signalling. Consistent with this, TBXT expression is increased in YAP1 knockout human gastruloids induced by BMP4 stimulation ^64^. Moreover, active YAP was downregulated *in vivo* during primitive streak formation in TBXT expressing cells (**Figure 4h**).

The node has been hypothesized to act by protecting the prospective somite and neural territory from surrounding BMP signalling activity^38,65^ and our data suggest a highly specialized node architecture. We found that BMP and NODAL inhibition was critical to generate and maintain trunk progenitors and posterior identity. We demonstrate that unchecked WNT and FGF signalling promotes endoderm and lateral mesoderm differentiation via sequential production of endogenous NODAL and BMP. In part this explains why 3D gastruloid protocols that use CHIR and FGF contain substantial amounts of endoderm^7,10^. We exploit the signaling cascade to expand current models of human gastrulation with a preparation that results in the robust production of notochord cells surrounded by trunk tissue. Previously, notochord cells had been lacking from *in vitro* derived structures and notochord has proved difficult to generate by directed differentiation^66,67^. Our data suggest a specific timing and signalling regime are required. As a defining feature of the chordates, the notochord is responsible for patterning trunk tissues and consistent with this, the neural tissue adjacent to *in vitro* generated notochordal cells had the molecular characteristics of floor plate and ventral neural progenitors. Together the data provide new insight into the mechanisms organising the vertebrate body plan and provide a foundation for future synthetic tissue design.

---

**Fig.S1** - a, Integrated UMAP embedding of chick trunk (4-13S) coloured by the unbiased Leiden clustering used to form the final clustering of trunk populations on Fig.1b. b, Percentage of cells expressing different HOXB genes showing induction of HOXB9 between 7-10S.c, contribution of the individual stages to the main UMAP of Fig.1b and FigS1a. d, Force-directed graph embedding showing the main trunk populations presence in each individual stage. The plots highlight that in all stages the same populations can be found as expected for caudal elongation stages. e-g, UMAP showing the expression of key genes characterizing the PXM and PS; LPM and intermediate mesoderm; and notochord and endoderm populations found in the chick trunk (4S-13S).

**Fig.S2** - a-b, Integrated UMAP embedding of chick trunk (4-13S) showing the expression of several genes marking the surface ectoderm; the PreNeural, Neural and Neural crest populations present during chick trunk formation. c, Gene expression of anterior PreNeural genes identified by Trevers et al. (2021). d, UMAP embedding detail plots highlighting the expression of several genes at and around the NMP cluster. Scale with maximum counts is indicated by the squares.

**Fig.S3** - a, Integrated UMAP embedding of mouse trunk formation (E8.0-E8.5) from Chan et al. (2019). Gene expression of main markers for each trunk population. b, UMAP showing the expression of key genes characterizing the Primitive streak; Lateral Plate Mesoderm (LPM), notochord, NMPs, endoderm and intermediate mesoderm populations found in the mouse trunk. Unbiased leiden clusters used to define main trunk populations. c, Equivalent embedding of macaque trunk formation (CS9-CS11) from Zhai et al. (2022) coloured by the main trunk populations. Gene expression of main markers for each trunk population and the leiden clusters used to define them.

**Fig.S4** - a, Optical section of a 7S chicken embryo HCR stained for NEFM shows localization in the caudal portion of somites and in the mature neural plate border. b, Immunostaining of SOX1 highlighting the first SOX1+ cells in rostral part of the sinus rhomboidalis, immediately rostral and on the side of the median pit. c, HCR of a 7S chicken embryo stained for ADAMTS18, CYP26A1, TBX6 and CNTN2.

**Fig.S5** - a, Overview of the Nucleus pipeline: cascade R-CNN is used for object detection in each z-plane, 2D nuclei are aggregated in 3D nuclei models which are further refined by a correction step. b, (left table) Training datasets used to train and validate the neural network. (right) Histogram with the maximum nuclear lengths in each dataset. These can be used to guide image acquisition as to whether the network will perform as expected. (bottom) Performance table example showing how different architectures predict the validation set by different average precision metrics. c, Examples of 2D segmentations performed by this tool.

**Fig.S6** - a, 3D centroids given by the Nucleus pipeline where selected by choosing a rectangle (x-, y-position) at a given dorso-ventral location (z-position). Dorsal and ventral positions around the median pit where picked and overlayed on a DAPI image. HCR average cell levels of NOTO and CNTN2 are plotted in red for the cell cohorts chosen. b, Immunostaining showing TBXT and FOXA2 co-localization at the medial pit marked by simultaneous high NOTO and high CNTN2. c, Transcriptional similarity ordering (pseudotime analysis) of mouse single cell data showing gene expression of several lineage and signalling genes. Cells corresponding to PXM, NMPs and Neural tissue were taken from clusters 9,8,2,11 (Fig.S3a).

**Fig.S7** - a, Monolayer differentiation of NMPs as in Figure 3a stained for SOX2 and TBXT. Nuclear levels of both markers showing a broad range of levels. b, SOX2 and TBXT nuclear levels in day3 posterior neuruloids were discretized into low, mid and high-levels. Optical section showing high SOX2 cells located at the center of the colony whereas high TBXT cells occupy a position at the periphery. Mid TBXT+SOX2+ cells localize to an intermediate ring in the colony. c, UMAP plot of posterior neuruloid day3 single-cell transcriptomics showing additional gene markers for mesoderm population and SHH expression in the notochord progenitors.

**Fig.S8** - a, Immunostaining overview of a micropatterned coverslip stained for pERK1/2 of posterior neuruloids at 12 hours post-induction. b, Staining of pERK1/2 and TBXT induction with different FGF concentrations, MEK inhibition and with no WNT agonist (CHIR; just N2B27+FGF). c, Dependency of mesodermal populations (TBX6 and TBXT) in the day3 posterior neuruloid with various levels of FGF.

**Fig.S9** - a, Total YAP and active YAP staining of the posterior neuruloid at 12h post-induction. Treatment with TRULI results in striking accumulation of YAP in the nucleus. b, Further characterization of the TRULI-treated posterior neuruloid showing no TBX6 or FOXA2 cells at day3. c, Morphology of TRULI-treated posterior neuruloid at day3 highlighting lower TBXT induction at the top and center of the colonies with only a few edge cells staining positive. d, additional stains of posterior neuruloid at 12h post-induction without and with TRULI showing a dampened pERK1/2 ring and no TBXT+ cells as well as the nuclear accumulation of YAP. Differences in phalloidin staining were also observed.

**Fig.S10** - a, Cross-species gene expression comparison of the notochord cell populations in chick, macaque and mouse single-cell transcriptomics data. b, Direct-force graph embedding layout with the integrated transcriptomic data of posterior neuruloid together with the 24h and 48h delay in TGFb inhibitors protocol variations. Gene expression of additional lateral plate mesoderm marker genes. c, Confusion matrix for the random forest classifier trained on chick and mouse cell clusters identified during trunk formation. A high recovery is observed for each cell population and the overall accuracy is 72% (with 25% data hold for testing). 166 genes representing the intersection of highly variable genes across species were used to train the classifier. d, Random forest classifier predictions with macaque and human cell queries. Macaque cell populations were used to independently validate the classifier with different in vivo data before comparing with human cells from posterior neuruloids. The different clusters were obtained from the combined cells across all BMP and NODAL inhibition delay treatments, e.g. the endoderm cell cluster is only present in the 48h treatment whilst notochord cells are mostly present in the 24h delay treatment.

**Fig.S11** - a, Protocol schematic of 24h delay 3D gastruloid cultures. b, Phase-contrast images of two juxtaposed 96-well round bottom plates. c, Two gastruloids, one positive for a TBXT streak and one negative. d, Automatic U-net segmentation of phase-contraste images with quantification of area versus solidity (area/ convex area) show the spread in morphology obtained in these preparations. e, Single-z plane of the inside of a gastruloid showing the presence of FOXC2+ somitic mesoderm in these 3D preparations. f, Optical section of gastruloid stained with the ventral neural markers NKX6.1 and OLIG2 around the TBXT+ notochord streak. f, Immunostaining of FOXA2, OLIG2 and SOX2 showing the presence of floor plate cells and the appearance of OLIG2+ cells.

## Methods

### Single-cell chick transcriptomics

Fertilized hens’ eggs obtained from Henry Stewart & Co. Ltd were incubated for 29 to 45 hours at 38°C with ~40% humidity to yield a minimum of 4 embryos with specific somite numbers: 4, 7, 10 and 13somites. All embryos were dissected to retain tissue caudal to the third somite (inclusive). A single-cell suspension was obtained by incubating the dissected embryos at 37°C in a dissociation solution consisting of accutase (Stemcell Technologies) with 3U/mg Papain (Sigma-Aldrich, 10108014001) and 1mg/mL of Collagenase 4 (Gibco, 17104019) for 20 min. Half-way through incubation and at the end, the embryos were mechanically dissociated with a P1000 pipet. After dissociation 200uL of resuspension buffer (DMEM/F12 with 1% BSA) was added. The cell suspension was then spun for 4min at 0.6xg, resuspended in 250μL of resuspension buffer and filtered through a 40μm Flowmi cell strainer (cat. no. 136800040) and twice through a pre-wet 20μm pluriSelect strainer (43-10020-60). The yield and cell viability for the 4S, 7S, 10S and 13S samples were 640, 500, 1300 and 1600 cells/μL with a viability of 88%, 93%, 92% and 95% respectively. A single-cell suspension was loaded independently for each sample onto the channels of Chromium Chip G for use in the 10x Chromium Controller (PN-1000120) with the goal of obtaining 10000 cells. The cells were partitioned into nanolitre scale gel beads in emulsions (GEMs) and lysed using the 10x Genomics Single Cell 3’ Chip V3.1 GEM, Library and Gel Bead Kit (PN-1000121). cDNA synthesis and library construction were performed as per the manufacturer’s protocol for the Chromium Single-Cell 3’ mRNA V3.1 protocol. cDNA amplification involved 12 PCR cycles. Libraries for the samples were multiplexed so that the number of reads matched one lane per sample and sequenced on an Illumina HiSeq4000 using 100 bp paired-end runs.

### Chick single-cell RNA-seq analysis

Reads were aligned to the *Gallus gallus* GRCg6a.101 reference genome using CellRanger (v4.0.0, 10X Genomics) and a custom-made reference 10X package including a gtf file with the protein coding, pseudogene and lncRNA gene biotypes. Read counts were computed using DropEst^68^ (v0.8.6) with the parameters “-f -V -w -L eiEIBA”. The remaining analyses were performed using Scanpy^69^ (v1.7.0) unless otherwise indicated. Data for the different chick stages were independently filtered for high percentages of mitochondrial UMIs (6-7.5%) and low total counts (2500). Potential doublet cells were filtered out using Scrublet^70^ with thresholds between 0.2-0.3. Counts were normalized to a target sum of 10000 excluding 1% of highly expressed genes. “Highly variable genes” were called using Scanpy’s function with default parameters. Data of different stages were integrated using harmonypy, a python port of the harmony^71^ R package by Ilya Korsunsky. Clustering of cells was performed unbiasedly using the Leiden algorithm with a resolution parameter of 3.5. PCA and UMAP embedding were run with default parameters, with neighbourhood graph computed for 10 neighbours and 40 principal components. Pseudotime inference was performed in R using Slingshot^36^ for the clusters between PXM and Neural.

### Chick RNA fluorescence in situ hybridization

Third-generation in situ hybridization chain reaction (HCR) DNA probe sets for the chicken mRNA genes NOTO, TBX6, CNTN2, ADAMTS18, NEFM and CYP26A1 together with HCR amplifiers, HCR probe hybridization buffer, and HCR probe wash buffer were ordered from Molecular Instruments^34^. Chicken embryos with 7 somites were dissected as to preserve caudal tissue and multiplex in situ hybridizations were performed according to the manufacturer’s protocol (Molecular Instruments HCR v3.0rev7 protocol for whole-mount chicken embryos). Embryos were mounted in ProLong SlowFade Mountant (Invitrogen, S36917) and imaged on a Leica SP8 confocal microscope.

### Nucleus segmentation pipeline

To segment 3D nuclei in whole-mount HCR chick embryos confocal stacks and posterior neuruloid micropatterned colonies, we developed a bespoke pipeline using Detectron2^72^, a pytorch-based computer vision library. To train the model, we manually segmented 13 cropped images of micropatterned neuruloids with a total of 928 nuclei instances. With these images we used transfer learning from an ImageNet pre-trained cascade R-CNN architecture^73^ to obtain an AP@[0.5:0.95] (average precision) of 55% and an AP0.5 of 86% on the validation set. The resulting 2D segmentation forms the basis for 3D consolidation. To merge nuclei detected in each z-plane of the confocal stack, we employ a supervised strategy. First, we construct a graph linking nuclear masks across z-planes if they share a minimum area overlap of 30%. This results in a collection of connected components (subgraphs where nodes are nuclei linked across z-planes) that are further refined via two manually defined thresholds, one with the typical number of z-planes for a single nucleus and another with a hard, upperlimit of this value. Given a particular subgraph with a number of nodes greater than the defined thresholds, we sequentially prune edges in the following order: first for edges with Jaccard distance greater than 0.7, then the edge with highest Jaccard distance for nodes sharing the same z, then for the next edge with the highest distance and finally, if the subgraph is still larger than the hard limit and no distance is above 0.2, a random edge is taken. Each individual subgraph constitutes a single-nucleus model. All nuclei are taken to generate a 3D mask that is used to calculate nuclear features such as average channel intensity.

### Human embryonic stem cell culture

Human ESCs from H9 line (WiCell) were routinely cultured in StemFlex medium (Thermo Fisher Scientific A3349401) on 0.5-mg/cm2 laminin-coated plates (Thermo Fisher Scientific A29249). Cells were passaged using ReLeSR according to the manufacturer instructions (StemCell Technologies #05872). Cells were tested for *Mycoplasma spp*. at 3-month intervals.

### Generation of micropatterned posterior neuruloids

Coverslips with micropatterned laminin were generated using a protocol adapted from ^74^. Briefly, isopronanol-cleaned 18mm coverslips were UVO-cleaned for 10min prior to incubation with PLL-g-PEG(5) for 1 hour in the dark. Coverslips were rinsed 3 times in deionised water before placing them on a custom-made chrome mask previously activated by UVO-cleaning (2 min). Close contact between mask and coverslip was ensure by pressing firmly with a pipette tip. The reverse, silver-side of the mask was exposed to UV during 8min after which coverslips were put in 70% Ethanol during 15 min. The patterned coverslips were let to dry and used within 4 weeks.

To seed cells, coverslips were first incubated for 3h at 37°C with rh-Laminin-521 (cat.no. A29248) diluted 1:10 in PBS +/+ (cat. no. 14040-091) and thoroughly washed with PBS+/+ as detailed in ^46^. Cells were then dissociated by washing with PBS-/- once followed by incubation with accutase for 5min at 37°C. Mechanically dissociated cells in accutase were diluted with 4x volume of StemFlex with 10uM Y-27632 (ROCK inhibitor, Tocris, #1254) and manually counted with Trypan Blue. Concentration of the single-cell suspension was adjusted to 670,000 cells/ mL with StemFlex with 10uM Y-27632. Cells were seeded by adding 3 mL of the single-cell suspension to the coated coverslip in a 6well plate well. After 3 hours the wells with coverslips were washed once with PBS-/- and the media replaced with fresh StemFlex for overnight incubation. Next day (18h after), the colonies were induced by first washing with PBS-/- then adding 2.5mL of induction media consisting of N2B27 media with CHIR99021 (3uM), FGF2 (5ng/mL), SB431542 (10uM) and LDN193189 (0.1uM). The media was replaced next day. The coverslips were fixed at day 3 (80 hours) in 4% fresh PFA for 30min at room temperature, washed 2x in PB S-/- and stored at 4°C until further analysis.

### Single-cell transcriptomics of posterior neuruloids

Coverslips with micropatterned posterior neuruloids and the two protocol variations of 24h and 48h delay in the TGFβ family NODAL and BMP inhibitors (SB431542 and LDN193189) were dissociated by first washing with PBS-/- and then incubating with accutase for 10min at 37°C following mechanical dissociation. Accutase suspension was then diluted in 4x volume with resuspension buffer, washed and strained as described above for chicken transcriptomics. The yield was 1110, 950 and 1170 cells/uL respectively with a viability >95%. The samples were separately loaded for capture with the Chromium System using the Single Cell 3’ v3.1 reagents (10X Genomics). Reads were aligned to the *Homo sapiens* GRCh38-3.0.0 reference genome using CellRanger (v4.0.0, 10X Genomics) and the analyses were performed using Scanpy^69^ (v1.7.0). Cells were filtered for a minimum of 200 genes, mitochondrial UMIs between 1-20%, total counts between 10,000-50,000 and doublet cells were filtered using Scrublet^70^ with a threshold of 0.2. Counts were normalized to a target sum of 10000 excluding highly expressed genes. “Highly variable genes” were called using Scanpy’s function with default parameters. Data of different conditions were integrated using harmonypy^71^. PCA and UMAP were run with default parameters, with neighbourhood graph computed for 15 neighbours and 30 principal components. A 2D embedding using force-directed graph drawing was computed by taking the UMAP coordinates as initial position.

### qPCR analysis of micropatterns

Plates with the micropatterned coverslips were washed with PBS-/- and the cell colonies lysed in RLT buffer (QIAGEN 1015762). RNA extraction was performed using RNeasy mini kit (QIAGEN 74106) according to the manufacturer’s instructions. cDNA synthesis was performed using Superscript III (ThermoFisher 18080051) from 1ug RNA using random hexamers and was amplified using PowerUp SYBR green (Applied Biosystems A25918). qRT-PCR was performed using the QuantStudio 12K Flex Real-Time PCR system (ThermoFisher) and the SYBR Green PCR assay (ThermoFisher A25742). Expression values for each gene were normalized against ATPF1, using the delta-delta CT method implemented in the ‘pcr’ R package. Relative expression across the time-course was computed by normalizing log10 expression+pseudo-count to the maximum value. Primer sequences are given in Table S2.

### Three dimensional cultures

To generate 3D cultures we first started a pre-culture of human ESCs by seeding 200,000 cells onto a 6cm petri dish with StemFlex media with 10 uM Y-27632 (ROCK inhibitor) for 2 days. Cells were then dissociated using 0.5mL of accutase for 4 min and added to 4mL of StemFlex with Y-27632 for manual counting with Trypan Blue. A total of 600 cells were seeded on each 96-well plate well with 40uL of StemFlex with Y-27632 and the plate was spun for 3 min at 300 rpm. The cells were then allowed to aggregate for 5 hours before slowly adding 150uL of StemFlex media. Cell aggregates were induced 19h after (next day) by replacing twice the media present with 150uL of 3N media with 3uM CHIR and 5ng/uL FGF2. After 24 hours the media was replaced with the 3N media used in the posterior neuruloid protocol which additionally included LDN and SB for two days. At day3, the media was replaced with 3N media supplemented with 40nM all-trans retinal (Sigma, CAS 116-31-4) for two days. At day 5, media was changed to 3N for two additional days. At the day 7 end-point, cultures were fixed in fresh 4% PFA for 1 hour at room temperature and thoroughly washed with PBS -/- before further analysis. Shape analysis of 3D cultures was performed from phase contrast microscope images segmented using a custom-made neural network pipeline employing the deep learning library fast.ai (v2.7.10). Briefly, 25 phase images were manually segmented, augmented using the albumentations package, and used to train a dynamic U-Net with *resnet34* architecture that includes self-attention layers and Mish activation function. The accuracy of this semantic segmentation task was 0.96.

### Immunostaining

Micropatterned cells and 3D colonies were blocked and permeabilized for 1 hour at room temperature in PBS-/- with 1% Triton-X, 10% DMSO, 10%SDS and 4% Normal Donkey Serum (D9663-10mL Sigma). They were then rinsed for 1 min in PBS-/- before overnight incubation with primary antibodies at 4°C. After incubation, samples were washed 3 times for 5min in PBS-/- and incubated overnight with secondary antibodies conjugated with Alexa Fluor 488, 555, 594 and 647 (1:1000 dilution) and 10 ng ml-1 of DAPI (ThermoFisher Scientific). Finally, samples were washed for 10min in PBS-/- before mounting with ProLong Glass Antifade Mountant (Invitrogen, P36980).

### Data and code availability

Single-cell RNA sequencing data have been deposited under accession numbers GSE223189 and GSE224404 for chick trunk and human micropatterns respectively. The scripts to analyse single-cell transcriptomic data are in https://github.com/tiagu/trunk_dev_scRNAseq. The Nucleus segmentation pipeline is available at https://github.com/tiagu/Nucleus. The pipeline to analyse 3D cultures is at https://github.com/tiagu/gastrunet.

## Acknowledgements

We thank Nic Tapon, Tom Frith and Rory Maizels for constructive discussions. We are grateful to Naomi Moris, Slivia Santos and Aryeh Warmflash for critically reading of the manuscript. We thank Rocco D’Antuono and Donald Bell from the Advanced Light Microscopy STP, and Christina Dix from the Making Lab STP for their help. We all also grateful to the Advanced Sequencing STP, and to Christelle Soudy from the Chemical Biology STP and Birgit Aerne for TRULI.

## Funding

This work was supported by the Francis Crick Institute which receives its core funding from Cancer Research UK (CC001051), the UK Medical Research Council (CC001051), and the Wellcome Trust (CC001051); by the European Research Council under European Union (EU) Horizon 2020 research and innovation program grant 742138 and by the Wellcome Trust (220379/D/20/Z). For the purpose of Open Access, the author has applied a CC BY public copyright licence to any Author Accepted Manuscript version arising from this submission.

## Contributions

T.R. and J.B. conceived the project, interpreted the data, and wrote the manuscript. T.R. and A.L. dissected chick embryos and performed single-cell RNA-seq and HCR. T.R. performed human ESC experiments. T.R. devised and performed bioinformatic and image analysis. T.R. and M.D. characterized posterior neuruloids by qPCR.

## Notes

### Competing Interest Statement

The authors have declared no competing interest.

