## Supplementary Data for "Notochord and axial progenitor generation by timely BMP and NODAL inhibition during vertebrate trunk formation"

**Fig.S11** - a, Protocol schematic of 24h delay 3D gastruloid cultures. b, Phase-contrast images of two juxtaposed 96-well round bottom plates. c, Two gastruloids, one positive for a TBXT streak and one negative. d, Automatic U-net segmentation of phase-contraste images with quantification of area versus solidity (area/ convex area) show the spread in morphology obtained in these preparations. e, Single-z plane of the inside of a gastruloid showing the presence of FOXC2+ somitic mesoderm in these 3D preparations. f, Optical section of gastruloid stained with the ventral neural markers NKX6.1 and OLIG2 around the TBXT+ notochord streak. f, Immunostaining of FOXA2, OLIG2 and SOX2 showing the presence of floor plate cells and the appearance of OLIG2+ cells.

**Figure S1**

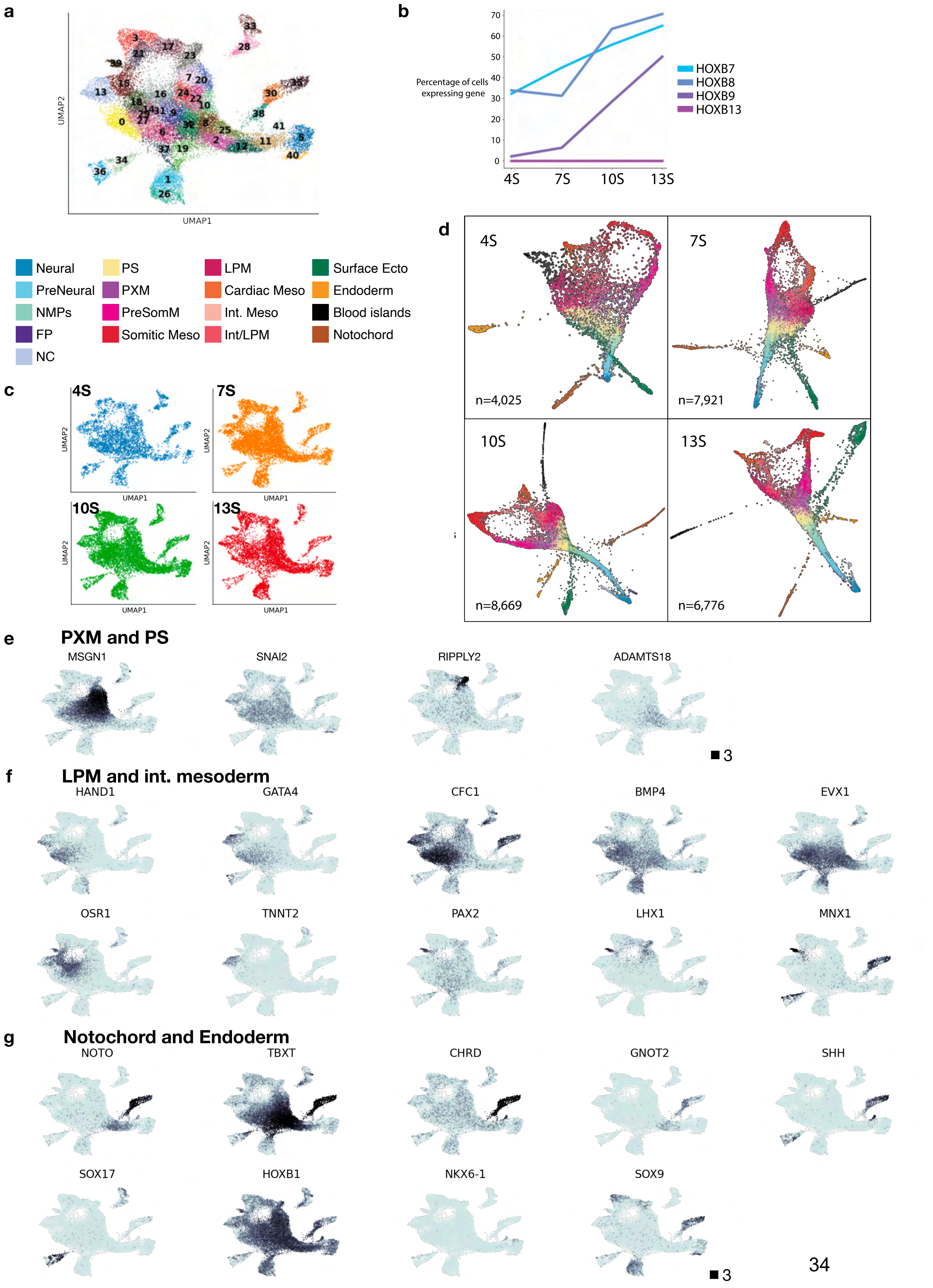
a
UMAP2
UMAP1
b
Percentage of cells expressing gene
4S
7S
10S
13S
HOXB7
HOXB8
HOXB9
HOXB13
Neural
PreNeural
NMPs
FP
NC
PS
PXM
PreSomM
Somitic Meso
LPM
Cardiac Meso
Int. Meso
Int/LPM
Surface Ecto
Endoderm
Blood islands
Notochord
c
4S
7S
10S
13S
d
n=4,025
n=7,921
n=8,669
n=6,776
e
PXM and PS
MSGN1
SNAI2
RIPPLY2
ADAMTS18
3
f
LPM and int. mesoderm
HAND1
GATA4
CFC1
BMP4
EVX1
OSR1
TNNT2
PAX2
LHX1
MNX1
g
Notochord and Endoderm
NOTO
TBXT
CHRD
GNOT2
SHH
SOX17
HOXB1
NKX6-1
SOX9
3

### Figure S2

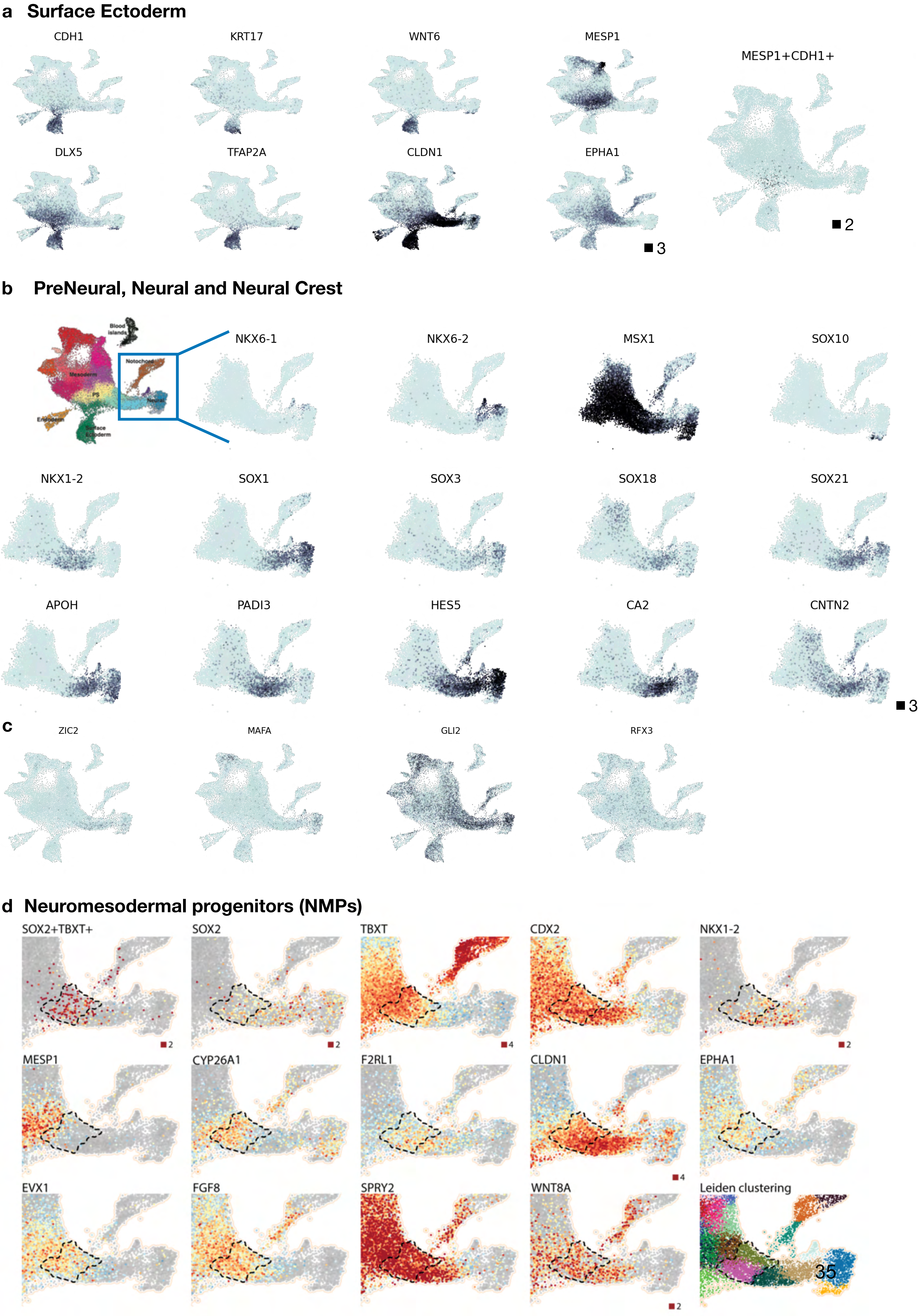
a Surface Ectoderm
CDH1
KRT17
WNT6
MESP1
MESP1+CDH1+
DLX5
TFAP2A
CLDN1
EPHA1
3
2
b PreNeural, Neural and Neural Crest
Blood islands
Notochord
Mesoderm
PS
Neural
Endoderm
Surface Ectoderm
NKX6-1
NKX6-2
MSX1
SOX10
NKX1-2
SOX1
SOX3
SOX18
SOX21
APOH
PADI3
HES5
CA2
CNTN2
3
c
ZIC2
MAFA
GLI2
RFX3
d Neuromesodermal progenitors (NMPs)
SOX2+TBXT+
SOX2
TBXT
CDX2
NKX1-2
2
2
4
2
MESP1
CYP26A1
F2RL1
CLDN1
EPHA1
4
EVX1
FGF8
SPRY2
WNT8A
Leiden clustering
2

**Figure S3**

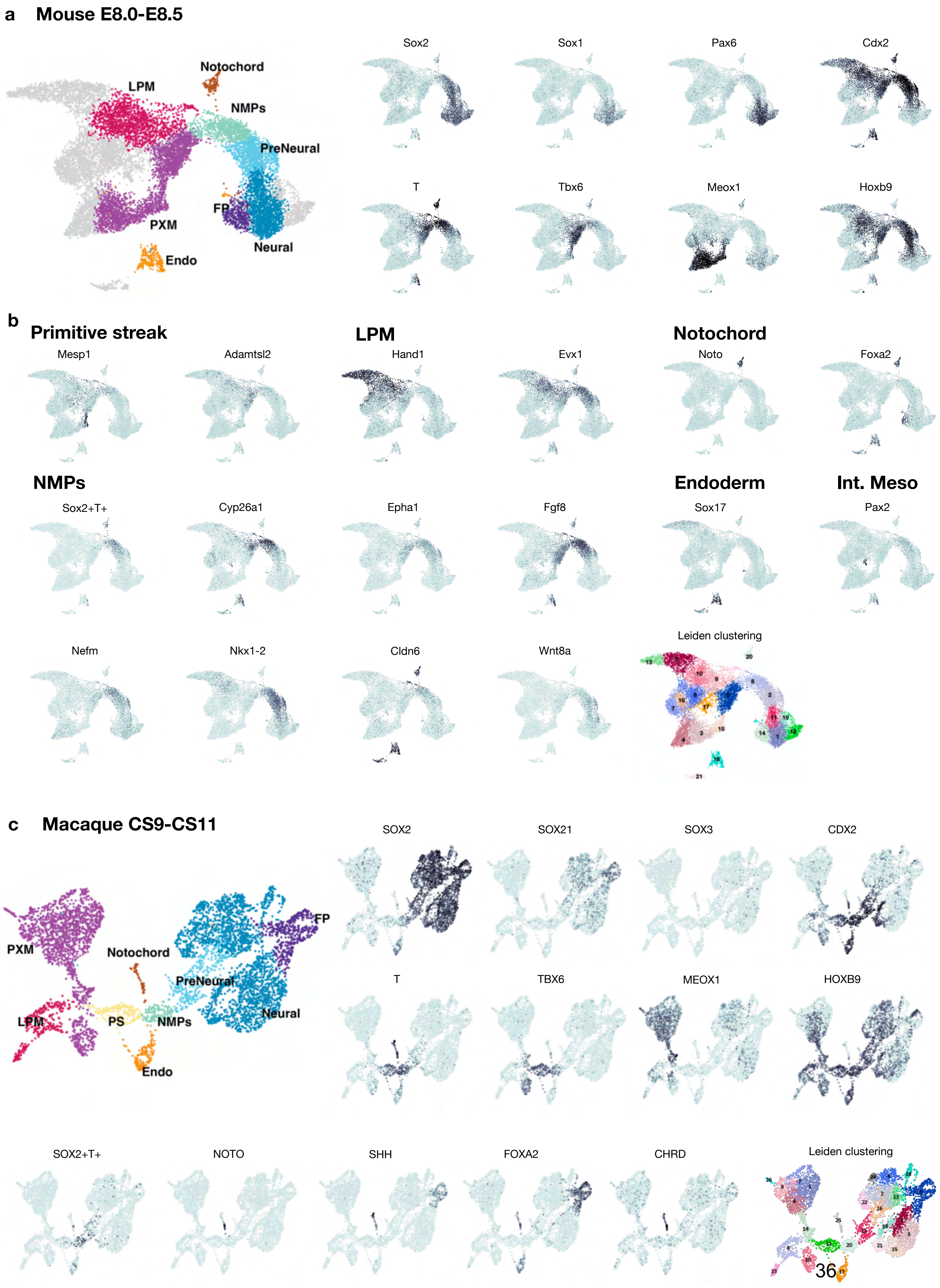
a
Mouse E8.0-E8.5
Notochord
LPM
NMPs
PreNeural
FP
PXM
Neural
Endo
Sox2
Sox1
Pax6
Cdx2
T
Tbx6
Meox1
Hoxb9
b
Primitive streak
Mesp1
Adamtsl2
LPM
Hand1
Evx1
Notochord
Noto
Foxa2
NMPs
Sox2+T+
Cyp26a1
Epha1
Fgf8
Endoderm
Sox17
Int. Meso
Pax2
Nefm
Nkx1-2
Cldn6
Wnt8a
Leiden clustering
c
Macaque CS9-CS11
FP
PXM
Notochord
PreNeural
LPM
PS
NMPs
Neural
Endo
SOX2
SOX21
SOX3
CDX2
T
TBX6
MEOX1
HOXB9
SOX2+T+
NOTO
SHH
FOXA2
CHRD
Leiden clustering

**Figure S4**

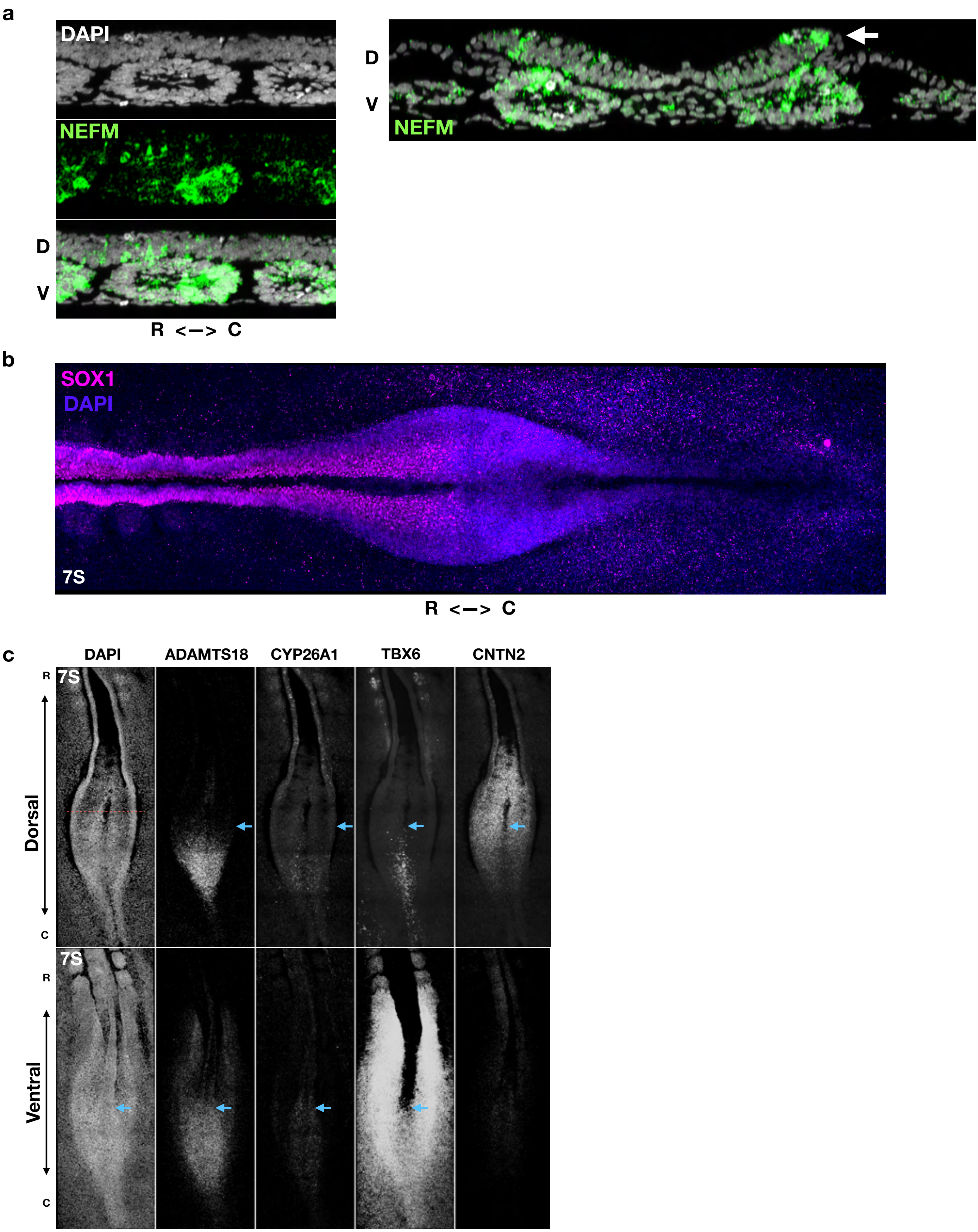
a
DAPI
NEFM
D
V
R <—> C
D
V
NEFM
b
SOX1
DAPI
7S
R <—> C
c
DAPI
ADAMTS18
CYP26A1
TBX6
CNTN2
R
7S
Dorsal
C
7S
R
Ventral
C

### Figure S5

#### a *Nucleus* pipeline for single-nuclei detection in dense colonies/ embryos

**1. High-confidence detection of 2D nuclei in single-z plane using CNN**

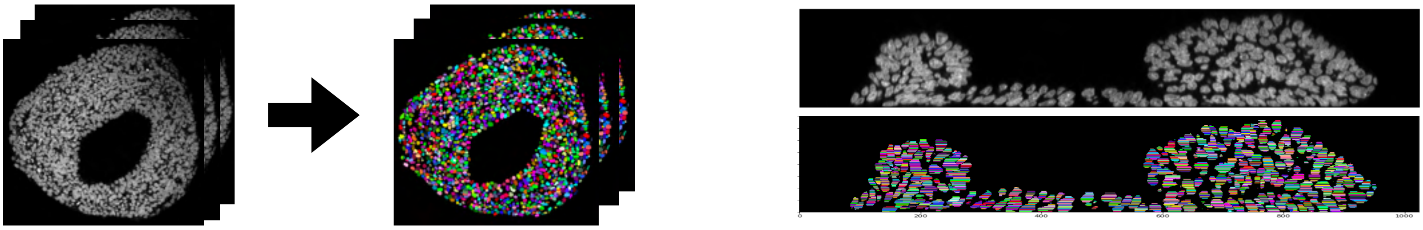

AP50 (avg precision at IoU.50) = 82-86%

**2. 3D consolidation and correction**

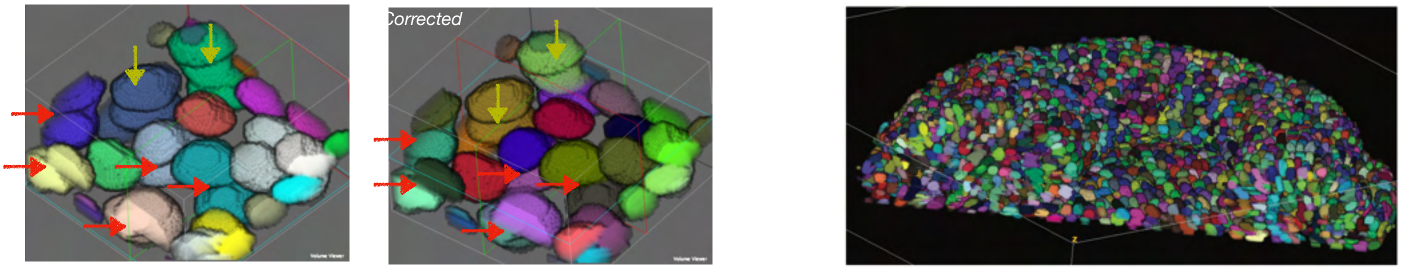


#### b Training set

| | #images | #instances | size | comments |
|---|---|---|---|---|
| nucleus_train | 6 | 221 | 256*256 | in vitro hESC assay * |
| nucleus_val (validation) | 4 | 141 | 256*256 | in vitro hESC assay * * |
| kromp_2019 | 52 | 1,704 | 640*512 | curated from Kromp et al. (2019) |
| segm_512 | 3 | 566 | 512*512 | in vitro hESC assay * |
| SC_human | 4 | X | 256*256 | Spinal Cord sections * |
| SC_mouse | 6 | X | 256*256 | Spinal Cord sections * |
| SC_sections (validation) | 5 | X | 256*256 | Spinal Cord sections * * |

* validation datasets
* new datasets manually segmented

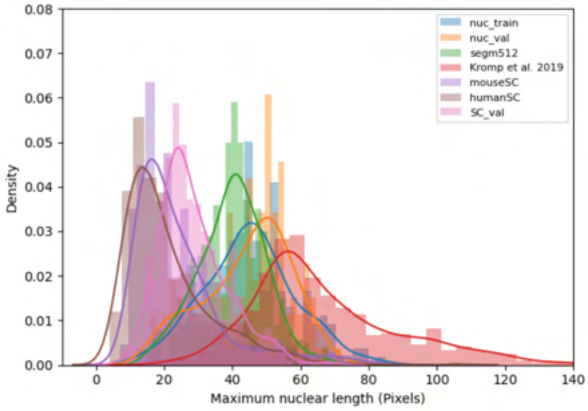


| | | | AP | AP50 | AP75 | APs | APm | AR |
|---|---|---|---|---|---|---|---|---|
| cascade_mask_rcnn_X_152 | SC_val | BBOX | 44.29 | 68.21 | 50.96 | 45.55 | 14.12 | 50.30 |
| | | SEGM | 42.59 | 67.31 | 47.68 | 42.72 | 40.00 | 48.60 |
| | nucleus_val | BBOX | 55.19 | 86.34 | 63.89 | 45.52 | 62.53 | 59.20 |
| | | SEGM | 53.11 | 86.55 | 61.16 | 43.87 | 60.02 | 57.40 |
| mask_rcnn_X_101 | SC_val | BBOX | 53.04 | 81.56 | 59.58 | 53.64 | 90.00 | 59.00 |
| | | SEGM | 53.89 | 82.52 | 61.08 | 53.83 | 90.00 | 59.10 |
| | nucleus_val | BBOX | 50.63 | 82.83 | 57.15 | 42.54 | 57.21 | 56.70 |
| | | SEGM | 49.54 | 86.29 | 54.32 | 42.65 | 54.92 | 55.00 |

#### c 2D segmentation examples

mouse hindbrain section

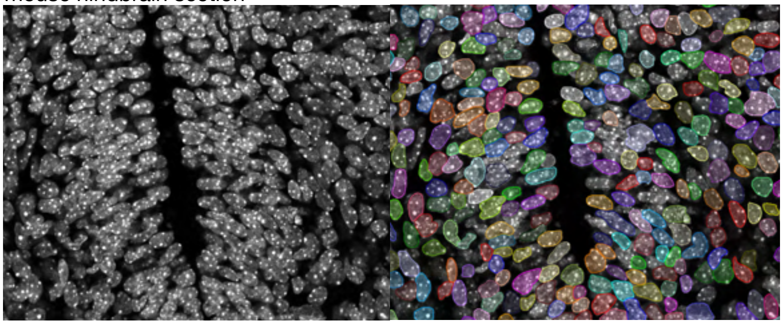

human spinal cord section

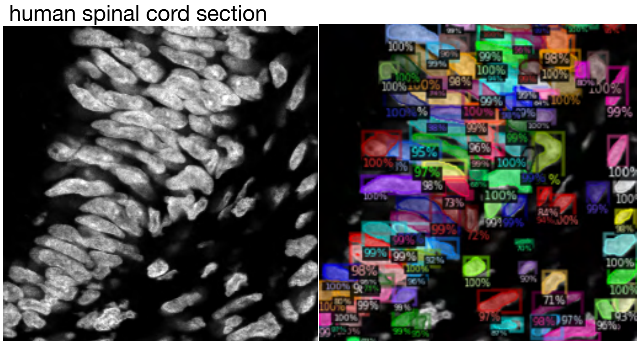

human in vitro organoid

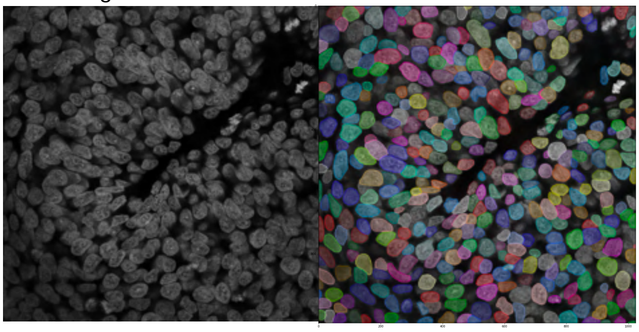

chicken embryo confocal optical section

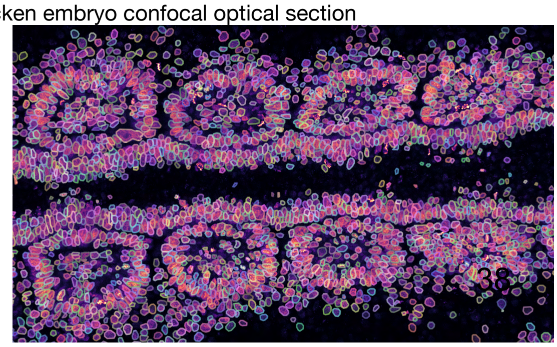

**Figure S6**

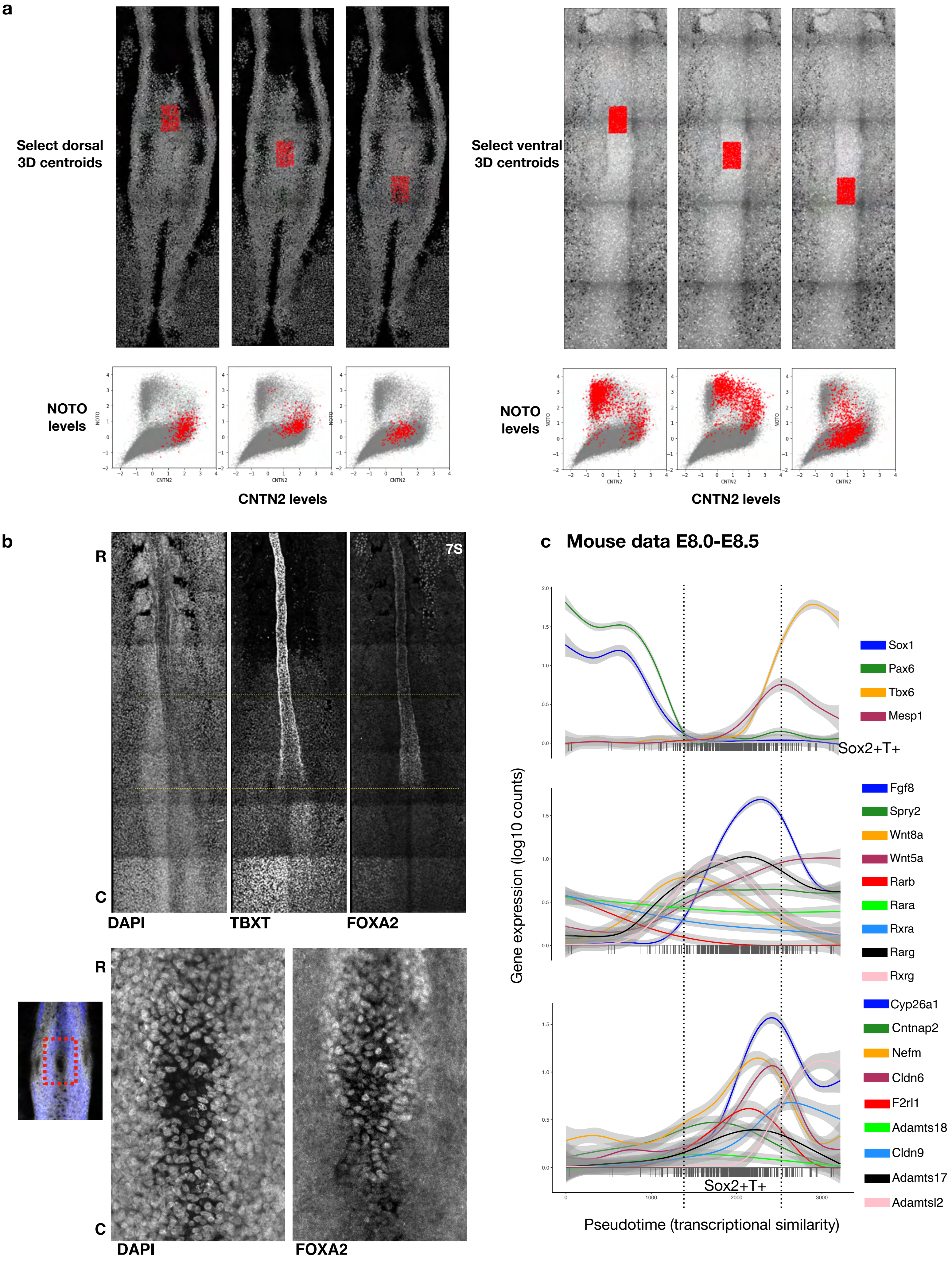
a
Select dorsal 3D centroids
Select ventral 3D centroids
NOTO levels
CNTN2 levels
NOTO levels
CNTN2 levels
b
R
C
7S
DAPI
TBXT
FOXA2
R
C
DAPI
FOXA2
c
Mouse data E8.0-E8.5
Gene expression (log10 counts)
Sox1
Pax6
Tbx6
Mesp1
Sox2+T+
Fgf8
Spry2
Wnt8a
Wnt5a
Rarb
Rara
Rxra
Rarg
Rxrg
Cyp26a1
Cntnap2
Nefm
Cldn6
F2rl1
Adamts18
Cldn9
Adamts17
Adamtsl2
Sox2+T+
Pseudotime (transcriptional similarity)

**Figure S7**

a

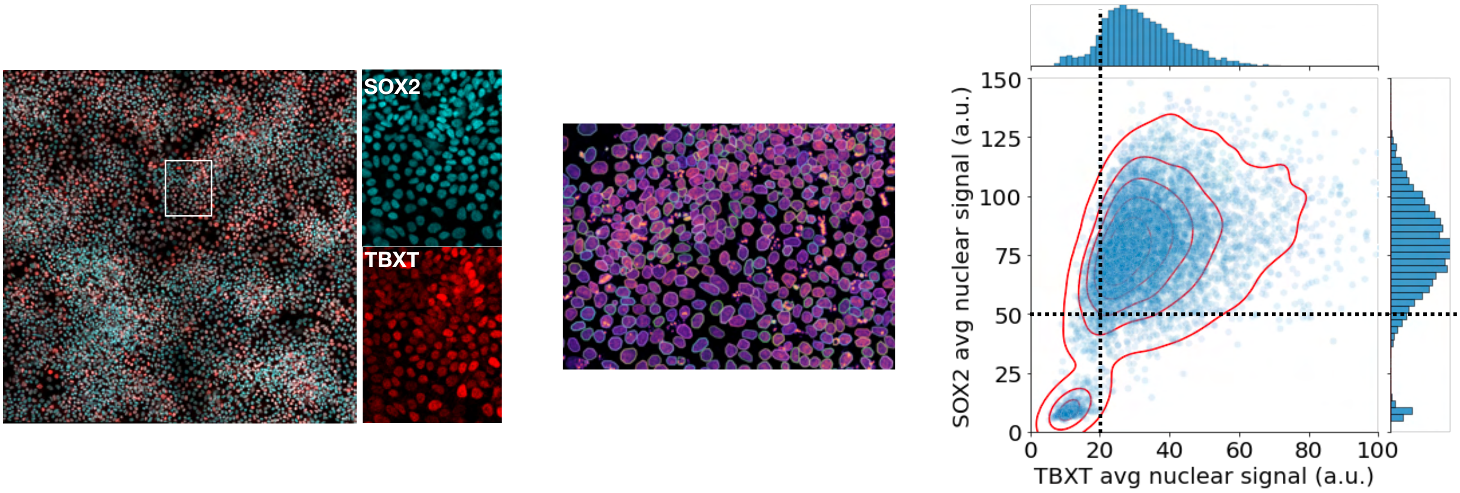


b

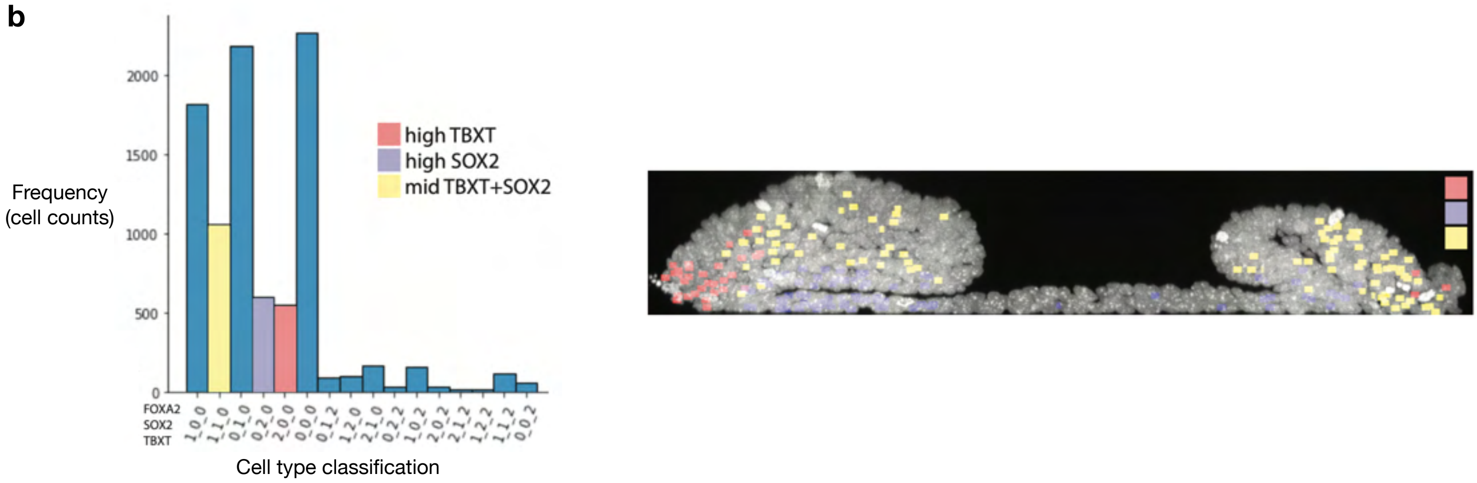


c

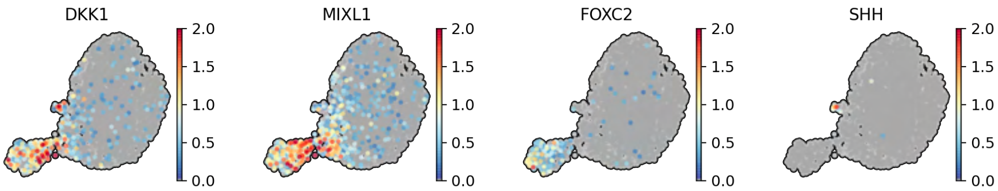

Figure S8

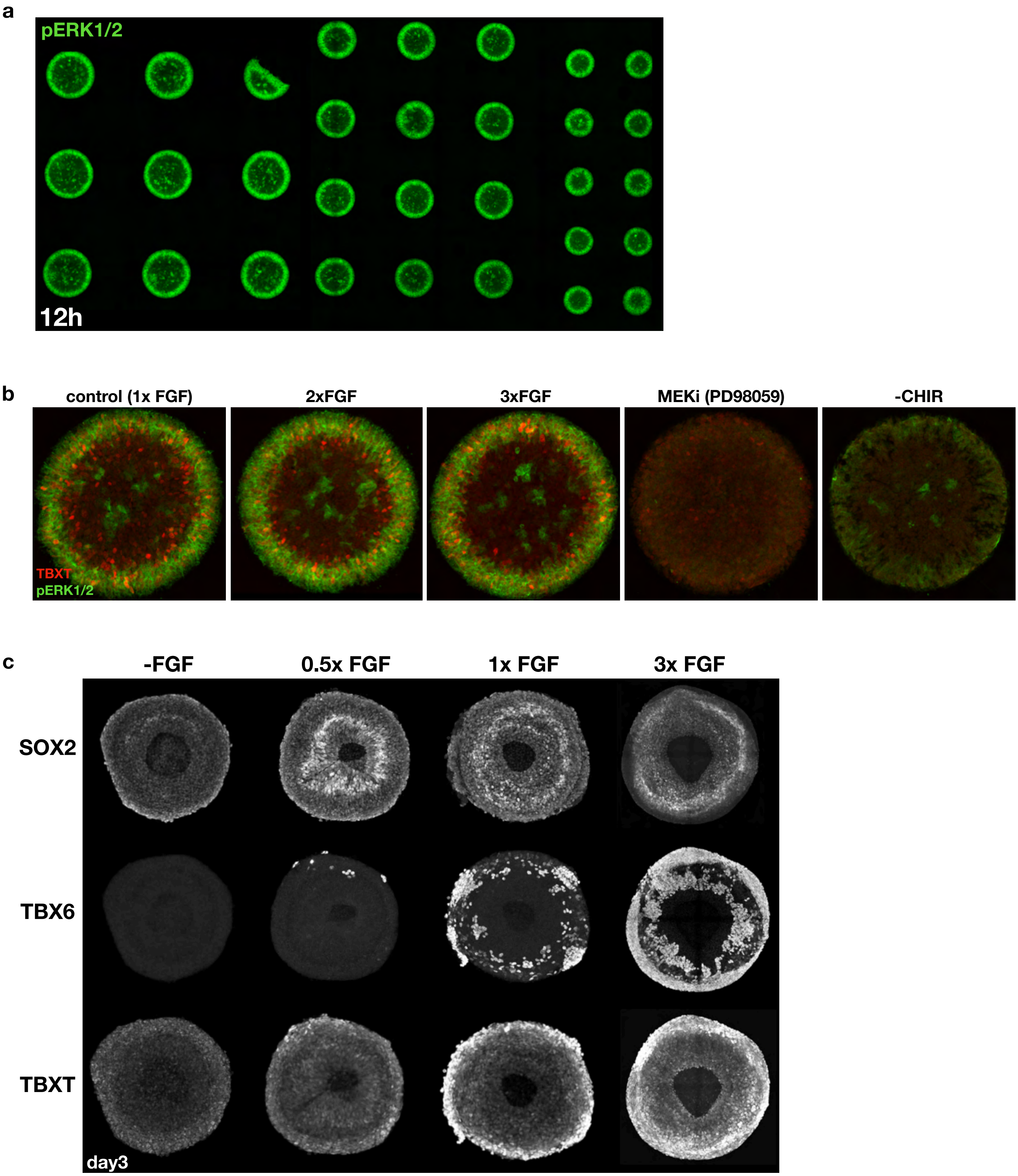
a
pERK1/2
12h
b
control (1x FGF)
2xFGF
3xFGF
MEKi (PD98059)
-CHIR
TBXT
pERK1/2
c
-FGF
0.5x FGF
1x FGF
3x FGF
SOX2
TBX6
TBXT
day3

Figure S9

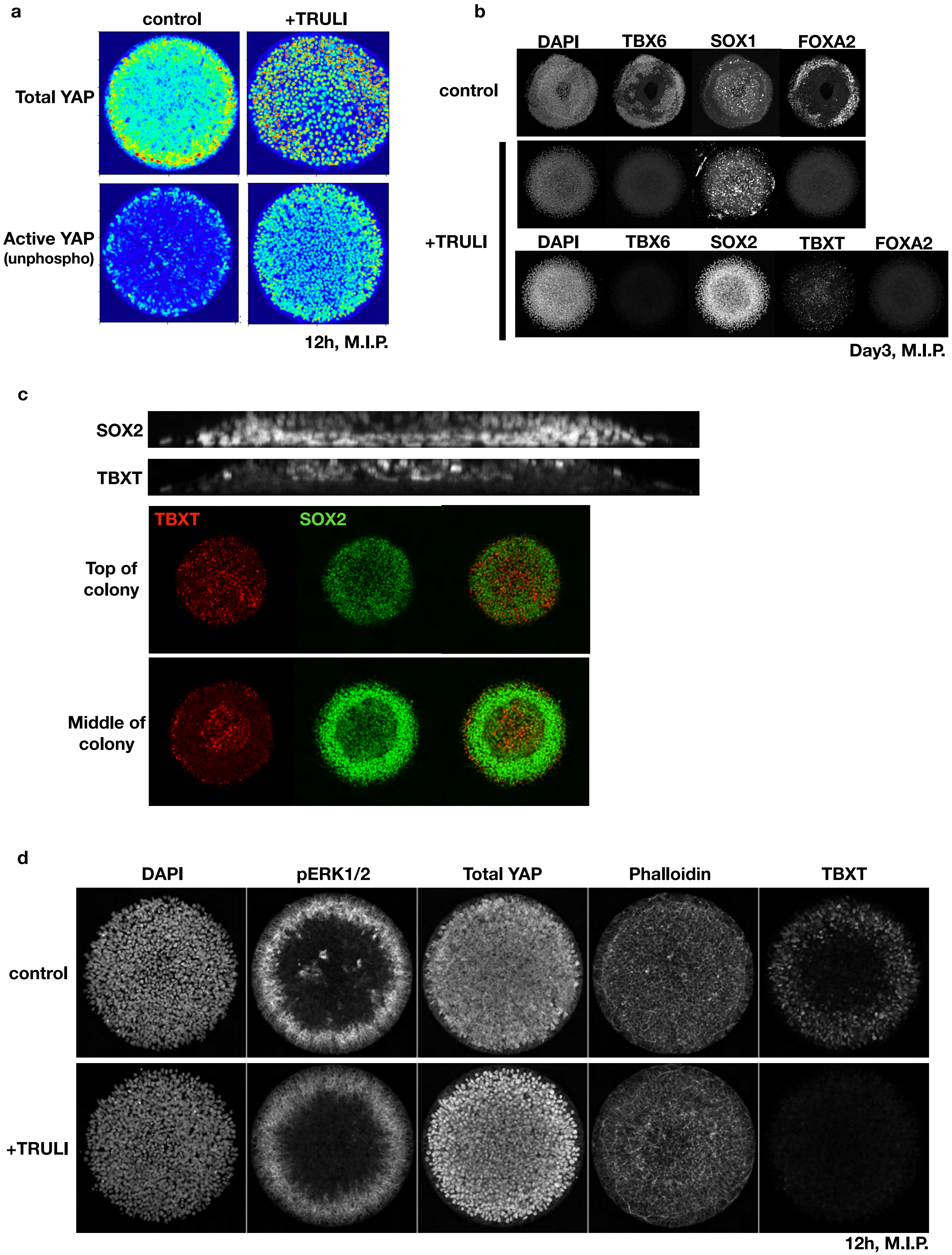
a
control
+TRULI
Total YAP
Active YAP
(unphospho)
12h, M.I.P.
b
DAPI
TBX6
SOX1
FOXA2
control
+TRULI
DAPI
TBX6
SOX2
TBXT
FOXA2
Day3, M.I.P.
c
SOX2
TBXT
TBXT
SOX2
Top of colony
Middle of colony
d
DAPI
pERK1/2
Total YAP
Phalloidin
TBXT
control
+TRULI
12h, M.I.P.

**Figure S10**

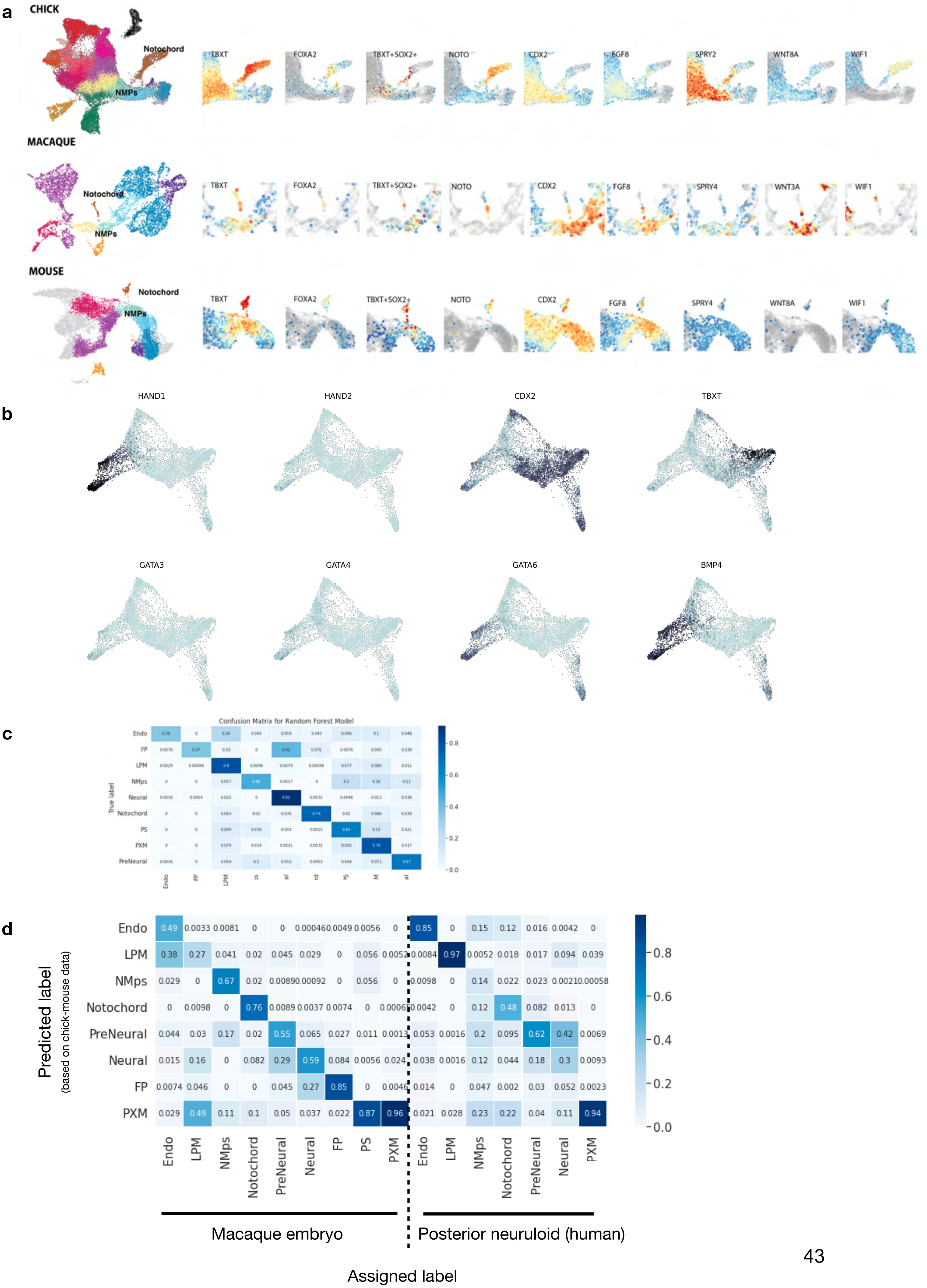
a
CHICK
Notochord
NMPs
TBXT
FOXA2
TBXT+SOX2+
NOTO
CDX2
FGF8
SPRY2
WNT8A
WIF1
MACAQUE
Notochord
NMPs
SPRY4
WNT3A
MOUSE
b
HAND1
HAND2
GATA3
GATA4
GATA6
BMP4
c
Confusion Matrix for Random Forest Model
True label
Endo
FP
LPM
NMps
Neural
Notochord
PS
PXM
PreNeural
d
Predicted label
(based on chick-mouse data)
Endo
LPM
NMps
Notochord
PreNeural
Neural
FP
PXM
PS
Macaque embryo
Posterior neuruloid (human)
Assigned label

**Figure S11**

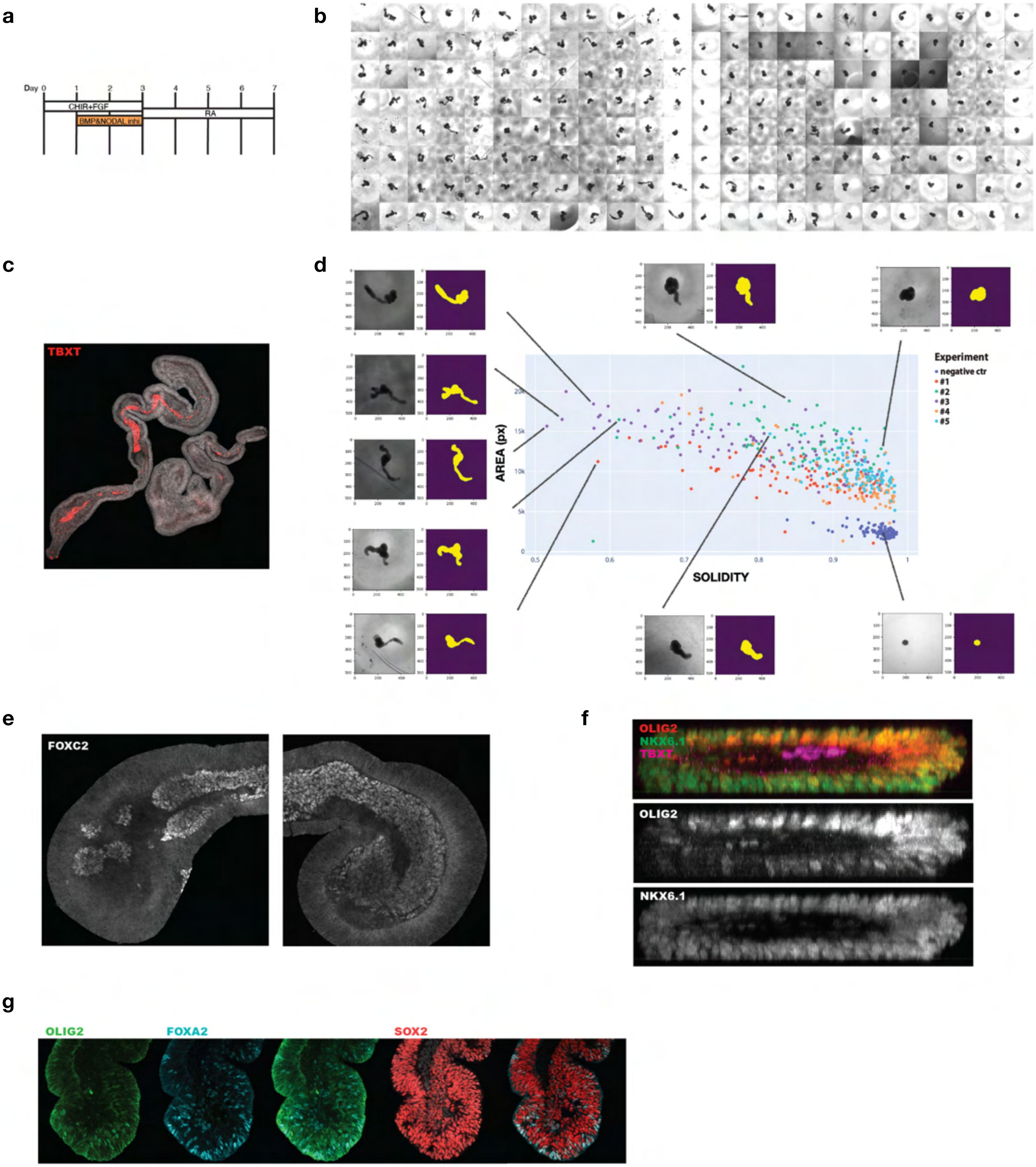
a
b
c
d
e
f
g
Day 0 1 2 3 4 5 6 7
CHIR+FGF
BMP&NODAL inhi
RA
TBXT
Experiment
negative ctr
#1
#2
#3
#4
#5
AREA (px)
SOLIDITY
FOXC2
OLIG2
NKX6.1
TBXT
OLIG2
NKX6.1
OLIG2
FOXA2
SOX2
